# Active learning of enhancer and silencer regulatory grammar in a developing neural tissue

**DOI:** 10.1101/2023.08.21.554146

**Authors:** Ryan Z. Friedman, Avinash Ramu, Sara Lichtarge, Yawei Wu, Lloyd Tripp, Daniel Lyon, Connie A. Myers, David M. Granas, Maria Gause, Joseph C. Corbo, Barak A. Cohen, Michael A. White

**Affiliations:** The Edison Family Center for Genome Sciences & Systems Biology, Washington University School of Medicine, Saint Louis, MO, 63110; Department of Genetics, Washington University School of Medicine, Saint Louis, MO, 63110; Department of Pathology and Immunology, Washington University School of Medicine, Saint Louis, MO, 63110

**Author notes:** Department of Genome Sciences, University of Washington, Seattle, WA, 98195.

## Abstract

*Cis*-regulatory DNA elements (CRE) are composed of transcription factor (TF) binding sites that direct cell type-specific gene expression. Deep learning is an emerging strategy to model CREs, but the genome offers too few training examples to learn the complex interactions between TF binding sites that govern CRE activities. We address this limitation using active learning to iteratively train models that predict enhancer and silencer activities in the developing mouse retina. Active learning doubled the performance of models trained on genomic data, resulting in models that accurately distinguish between enhancers and silencers composed of the same TF binding sites. The ability of these models to discriminate between functionally non-equivalent binding sites establishes active learning as an effective strategy for modeling regulatory DNA.

**One sentence summary:** Models of regulatory DNA learn critical interactions between transcription factor binding sites with active machine learning.

## INTRODUCTION

The “*cis*-regulatory grammar” of a cell type describes how the *cis*-regulatory activities of DNA sequences are related to the arrangements and interactions of the TF binding sites that they contain (*1*–*6*). The contribution made by a TF binding site to the activity of a CRE strongly depends on the context created by other features present in the local DNA sequence, especially other interacting TF binding sites. Because of context effects, different instances of binding sites for the same TF are often functionally non-equivalent, and CREs composed of similar TF binding sites often have dramatically different activities (*7*–*24*). Such functional differences between similar TF binding sites are why standard analyses of enriched TF binding motifs have limited power to distinguish genuine CREs from non-functional DNA, or to predict the effects of non-coding genetic variants on gene regulation. Models of *cis*-regulatory grammars that can accurately distinguish functionally non-equivalent TF binding sites would therefore address a major challenge in ongoing efforts to understand the role of the non-coding genome in human health and disease.

Machine learning presents an opportunity to train better models of *cis*-regulatory grammars, due to its power to discover predictive features in high dimensional data. Deep neural network models trained on large epigenomic datasets often predict TF binding and chromatin accessibility with high accuracy, and these models have revealed important contextual features of local DNA sequence that determine TF binding (*25*–*32*). However, models trained on massively parallel reporter gene assays (MPRAs) to predict CRE activity (*20, 33*–*40*) often perform less well than binding models, likely because the *cis*-regulatory grammars that govern activity depend on additional higher-order interactions between bound TFs and their associated co-factors (*2, 4*–*6, 41*). Such higher-order interactions may be responsible for the functional non-equivalence of identical instances of TF binding sites. However, because the number of possible arrangements of binding sites within CREs grows exponentially with the number of sites, a major obstacle to learning the effects of higher-order interactions is that the number of genomic examples of active CREs in any particular cell type is small relative to the scale of the training data needed to learn the interactions among binding sites (*42, 43*). As a consequence, current deep learning models of *cis*-regulatory activity typically uncover TF motifs with large independent effects on gene expression, which tend to be the same motifs identified by traditional motif-finding algorithms. To learn the more complex features of *cis*-regulatory grammars, additional training data can be generated by MPRAs that test synthetic DNA sequences. Yet the number of possible sequences vastly exceeds the number that can be feasibly synthesized and assayed. There is thus an urgent need for methods to prioritize informative training examples from the space of potential sequences, thereby leveraging the capacities of MPRAs and other functional genomics assays to generate large training datasets that are not limited by what the genome alone provides.

We address this problem by introducing a major modification to the current paradigm for learning *cis*-regulatory grammars. We deploy active machine learning (*44*–*46*) to iteratively train models on successive rounds of informative MPRA experiments. In contrast to current approaches that rely on a single round of training data, active learning offers a way to iteratively improve models by selecting new training examples based on their potential to improve the model. Active learning has been successfully applied to model metabolic networks (*47*), optimize cell culture media (*48*), perform *in silico* drug screens (*49*–*52*), identify TFs that drive cellular differentiation (*53*), select optimal training data for nanopore base calling (*54*), and design Perturb-seq experiments (*55*). Here we apply active learning to the problem of *cis*-regulation for the first time, using it to iteratively train models of a cell type-specific *cis*-regulatory grammar in the early postnatal mouse retina, an experimentally accessible portion of the developing central nervous system. The model more than doubled its performance relative to training on genomic examples alone, and it learned to distinguish functionally non-equivalent binding sites for multiple cell type-specific TFs.

## RESULTS

### Active learning applied to *cis*-regulation in the developing retina

We used active learning to iteratively train models of a cell type-specific *cis*-regulatory grammar in a developing neural tissue, the early postnatal mouse retina. DNA binding sites for the retina-specific TF Cone-rod homeobox (CRX) present a striking example of how sites for a single TF encode distinct activities in different CREs (*56*–*62*). CRX plays a central role in the terminal differentiation of several cell types of the neural retina, most notably photoreceptors, which derive from a population of common progenitor cells (*63*–*67*). CRX binds both enhancers and silencers (*13, 20, 68*), and it cooperates with the closely related homeodomain TF OTX2 and other lineage-specific TFs to both activate and repress different target genes (*10, 69*–*74*). CRX is a ‘K50’ homeodomain TF (i.e., it has a lysine (K) in position 50 of its homeobox) which binds a motif that is pervasive in the open chromatin of photoreceptors and bipolar cells (*60, 61*), two cell classes that together constitute nearly 90% of all cells in the mouse retina (*75*). Approximately 50% of open chromatin regions in rod photoreceptors contain a CRX motif, making it by far the most enriched cell type-specific TF binding motif in the most abundant cell type of the mouse retina (*60*). However, because CRX acts at both enhancers and silencers, the presence of a CRX motif is by itself insufficient to predict whether a CRX-bound DNA sequence will activate or repress transcription (*13, 16, 20, 61, 68, 76*). There are also many CRX-bound sequences that exhibit no *cis*-regulatory activity in the developing neural retina, similar to results reported for TF-bound sequences in cell culture models (*17, 77*–*80*). Thus a key challenge for models of *cis*-regulation is to learn how local sequence context encodes the non-equivalent functions of binding sites for multi-functional TFs such as CRX.

To learn complex interactions between TF binding sites that determine *cis*-regulatory activity in developing rod photoreceptors, we implemented an active learning strategy that leverages the capacities of MPRAs to measure tens of thousands of synthetic DNA sequences with designed perturbations (**Fig. 1a**). The core of the approach is to train models on successive rounds of new experimental data that include training examples that are *actively* selected for their potential to improve the model on the next training round. In each round, the model is trained on a cumulative dataset that includes data from all prior rounds, and performance is evaluated on independent test sets. New candidate sequences are then generated by performing millions of *in silico* perturbations to sequences in the current training data. The new sequence pool is filtered to remove any candidates that lack general sequence properties of potential photoreceptor *cis*-regulatory elements by scoring them with a Support Vector Machine (SVM) trained on measurements of open chromatin in rod photoreceptors (*76*) (**fig. S1**, Methods). The SVM classification does not depend on MPRA measurements. The filtered pool of candidate sequences is then sampled to obtain the next round of training examples by using the current model to pick sequences whose predicted activities are the most uncertain (Methods). Uncertainty-based sampling is the key step of active learning and is based on the premise that candidate sequences predicted by the model with the least confidence will be the training examples most likely to improve the model in the next round (*46*). Following uncertainty sampling, the selected sequences are synthesized and assayed, and the results are added to the cumulative training dataset. With this active learning strategy, models are iteratively improved beyond the performance achieved with only a single round of training data.

**Figure 1:**
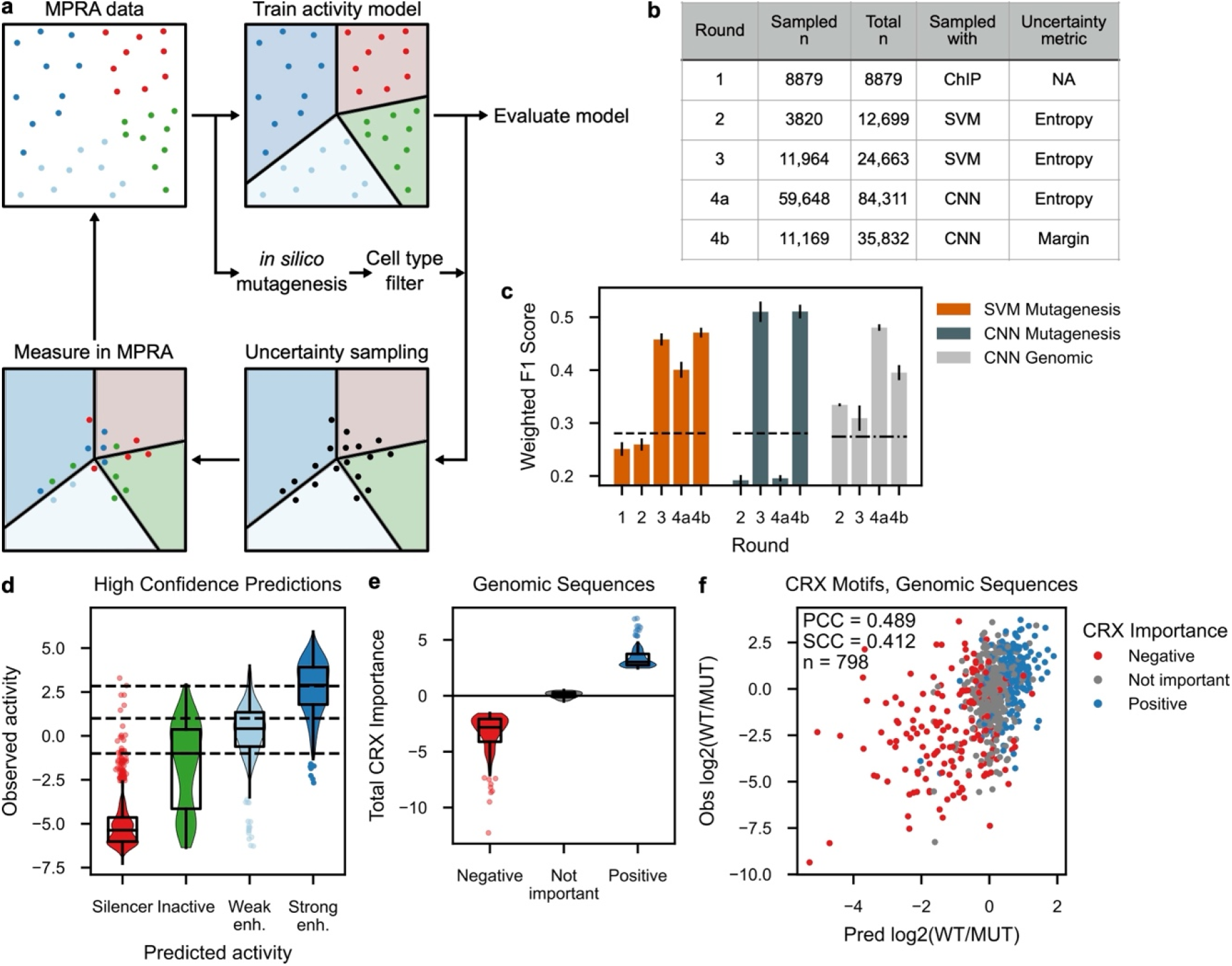
Iterative machine learning improves predictions of *cis*-regulatory activity. **a**, Summary of active learning approach. Colored dots represent sequences measured in MPRAs (dark blue, strong enhancer; light blue, weak enhancer; green, inactive; red, silencer), which are used to train a multi-class classifier (solid lines represent the margins between classifications inferred by the model and shaded areas correspond to the inferred activity classes). After generating a filtered pool of candidate sequences, those predicted with high uncertainty under the current model (black dots) are synthesized, measured by MPRA, and added to the training data for the next round of model fitting. **b**, Summary of the size and sampling method of each training data batch. **c**, Iterative improvement of model performance on the mutagenic series and genomic test sets. Horizontal lines represent accuracy expected from random guessing for the mutagenic series (dash) and genomic test set (dash-dot). Error bars denote one standard deviation based on ten-fold cross-validation of the newly added data. **d**, Observed activity of synthetic sequences predicted with high confidence, stratified by predicted activity class. Horizontal dashed lines correspond to cutoffs for the activity classes. Enh., enhancer. **e**, Stratification of genomic test set sequences by CRX importance. **f**, Predicted vs observed effect of mutating all CRX motifs in genomic sequences, colored by the total CRX importance. PCC, Pearson correlation coefficient; SCC, Spearman correlation coefficient.

To generate the data, we performed MPRAs in explanted mouse retina, a part of the central nervous system that is amenable to electroporation-based episomal reporter assays and which contains an abundant population of rod photoreceptors, which constitute nearly 80% of all cells in this tissue. We previously established MPRAs in retinal explants as a sensitive measure of *cis*-regulatory activity in differentiating rods (*13, 81*). We use the rod-specific *Rhodopsin* promoter as the basal promoter for the reporter gene libraries. The moderately high activity of the *Rhodopsin* promoter is a critical feature of this assay, because it allows us to measure both enhancer and silencer activity, ensuring that our training data include the full range of relevant *cis*-regulatory activities. We electroporate retinas at postnatal day 0 and harvest at postnatal day 8, a time window during which CRX actively promotes terminal differentiation of photoreceptors (*63, 64, 66*). We therefore assay CRX-targeted CREs in the cellular and developmental context in which they are natively active.

We trained initial models on a set of CRX-bound genomic sequences and then performed three rounds of active learning (**Fig. 1b**). We used two different four-way classifiers: a modified *k*-mer Support Vector Machine (SVM) and a convolutional neural network (CNN). An advantage of the SVM is that the feature encoding is predetermined, which allows it to be trained on smaller datasets. An advantage of the CNN is that it can flexibly encode higher order features, which may not be captured by the SVM. Each model predicts the probability that a given DNA sequence is a (1) strong enhancer, (2) weak enhancer, (3) inactive sequence, or (4) silencer in photoreceptors (Methods). Uncertainty measures are calculated from the probabilities assigned to each activity class (described below). We evaluated model performance on two independent MPRA test datasets that represent different prediction tasks. The first test set (“mutagenic series”) is an exhaustive motif perturbation analysis of 29 strong enhancers derived from the genome and each centered on a CRX binding motif (n = 711). This test set assesses the ability of the model to predict the effects of perturbations to TF binding sites in highly active enhancers. A limitation of this test set is that the classes are imbalanced (44% strong enhancers vs 3.7% silencers). The second test set (“genomic test set”) includes genomic CRX-bound sequences (*82*) that were not used elsewhere in the active learning pipeline (n = 1,723). The genomic test set reflects the natural ratio of enhancers to silencers among wild-type CRX-bound sequences (8% strong enhancers, 22% silencers) and sequences were not centered on CRX motifs. The SVM was only evaluated on the mutagenic series, because the SVM kernel function considers absolute *k*-mer positions, rather than relative positions, and thus requires input sequences centered on a CRX motif as a reference point.

### Active learning more than doubles model performance after exhausting genomic training data

Our initial SVM and CNN models (Round 1), were trained on an MPRA dataset derived from 164-bp photoreceptor-accessible chromatin sequences centered on a CRX motif (*20*). The dataset included 4,629 native genomic sequences, along with matched versions in which all CRX motifs in the native sequences were abolished with a point mutation known to abrogate binding. The SVM achieved a weighted F1 score (wF1) of 0.251 (**Fig. 1c, fig. S2**, and Methods), where a score of 0.28 represents random guessing on the mutagenic series and a score of one represents perfect four-way classification. The CNN did not generalize beyond the training data, presumably because the Round 1 genomic dataset was too small (**fig. S3**). Because most genomic CRX-bound sequences are in either the training dataset or the genomic test set, the performance of the initial models shows that there are not enough genomic examples of *bona fide* CREs to learn the context-dependent interactions between CRX motifs and other sequence features that specify non-equivalent functions of CRX motifs in enhancers, silencers, and inactive sequences.

After three rounds of active learning, the performance of these models more than doubled, on average (**Fig. 1c**, Rounds 2-4). In the final round, we tested two alternative sampling strategies (Rounds 4a and 4b). At Round 4b, the performance of the SVM increased to 0.471, while the performance of the CNN achieved a wF1 score of 0.511 on the mutagenic series and 0.395 on the genomic test set. Performance on the genomic test set was even higher at alternate Round 4a (wF1 = 0.480). These results demonstrate that iterative training with active learning substantially improves models of *cis*-regulation after exhausting genomic CRX-bound CREs as training and test data.

These global measures of model performance incorporate both the high- and low-confidence predictions. However, an advantage of our approach is that uncertainties are associated with each model prediction, and thus high-confidence predictions can be separated from low-confidence predictions. To test the accuracy of high-confidence CNN predictions, we synthesized and assayed 2,055 new synthetic sequences whose activity was predicted with high confidence. Remarkably, 72% of the high-confidence perturbations were correct (wF1 = 0.72, **Fig. 1d**). Furthermore, 14.5% of the high confidence predictions were off by only one class, indicating that the CNN learned the rank order of classes, even though this information was not explicitly provided. This shows that the internal confidence measures assigned by the models are reliable guides for the uncertainty sampling step of active learning. These confidence measures are also useful for downstream applications such as *de novo* enhancer design by focusing attention on the predictions that are most likely to be correct.

Discrete multi-class classifiers were used during active learning, but a model that predicts continuous values of CRE activity is more useful for making quantitative predictions about the role of individual TF binding sites. We trained a regression CNN on the cumulative training data of Round 4b and obtained a Pearson correlation coefficient (PCC) of 0.607 on the mutagenic series, and a PCC of 0.482 on the genomic test set (**fig. S4**). While model performance on the genomic test set is modest, this test set includes a large number of silencers, a class of CRE that current published models of MPRA activity do not predict at all.

### The model learned both enhancer and silencer effects of CRX motifs

Despite explaining only part of the variation in the test sets, the model achieved the major goal of distinguishing functionally non-equivalent CRX motifs in enhancers and silencers. We previously showed that experimentally mutating CRX motifs in enhancers tends to cause loss of activation, while mutating CRX motifs in silencers tends to cause loss of repression, though it was unclear why this happens (*20, 82*). The model now accurately predicts these directional effects. To test the model predictions, we used the model to assign importance scores to the CRX motifs in every sequence of the genomic test set (**Methods** and **fig. S5a**) and created three bins: CREs with negative CRX importance (most negative 10%), CREs with CRX importance near zero, and CREs with positive CRX importance (most positive 10%, **Fig. 1e**). When compared against experimental data of CRX motif mutations, CREs with the most negative CRX importance contained repressive CRX sites, while CREs with the most positive CRX importance contained activating CRX sites, confirming that the model correctly predicted the effects of CRX sites (**Fig. 1f** and **fig. S5b**). The model also correctly predicted non-functional CRX sites: CREs with CRX importance near zero exhibited only small changes in activity when CRX sites were mutated, despite those motifs being high scoring matches for the CRX position weight matrix. Unbinned predictions of CRX motif mutations made on the entire genomic test set achieved a similar performance (**fig. S5c**, PCC 0.421). Thus the model successfully learned the sequence context that distinguishes activating CRX sites in enhancers and repressive CRX sites in silencers, a crucial feature of the *cis*-regulatory grammar of developing photoreceptors that was not captured by the poorly performing initial models trained on genomic data alone.

### The model identifies causal differences between CREs with similar TF binding motifs

An important manifestation of how TF binding sites are often functionally non-equivalent is the finding that many TF-bound DNA sequences that reside in open chromatin nonetheless lack *cis*-regulatory activity (*13, 17, 77, 79, 83, 84*). Models of *cis*-regulatory grammar that correctly distinguish functionally non-equivalent sites should be able to identify the causal sequence differences between active and inactive sequences that contain similar TF binding motifs. To test the ability of our regression CNN to do this, we identified two rod-specific accessible chromatin sequences with one copy each of the CRX motif, and one copy each of a motif for the rod-specific TF NRL (**Fig. 2a**). One sequence was inactive, while the other was a strong enhancer when tested by MPRA. The sequences were centered on their respective CRX motifs. The NRL motifs were identical, although their position differed. A scan for other motifs known to be enriched in CRX-bound strong enhancers did not identify additional motifs in either sequence.

**Figure 2:**
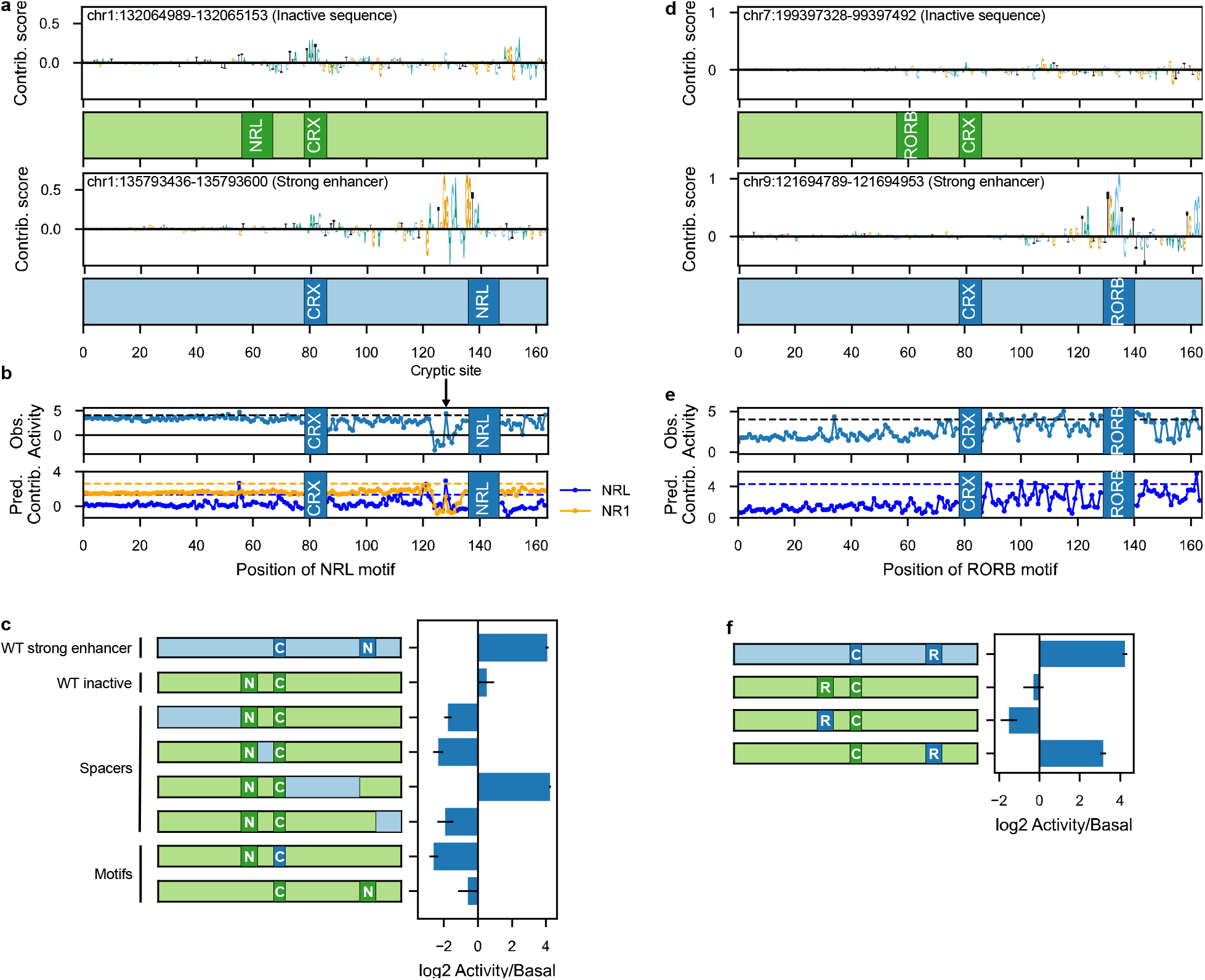
Regression CNN identifies causal sequence differences between active and inactive sequences. **a**, Relative nucleotide contribution scores and locations of known motifs for an inactive sequence (top) and a strong enhancer (bottom). **b**, Observed effect on activity of moving the NRL motif within the strong enhancer (top) and predicted contribution of the NRL and NR1-family motifs to activity (bottom). Horizontal dashed lines represent the activity (top) and motif contribution (bottom) of the wild-type sequence. **c**, Effect on activity of swapping strong enhancer regions into the inactive sequence. Cartoons show the chimeric constructs with colors matching (**a**). Error bars show standard deviation of one sequence across 3 replicates. C, CRX motif; N, NRL motif. **d**, Relative nucleotide contribution scores and locations of known motifs for a second inactive sequence (top) and strong enhancer (bottom). **e**, Observed effect on activity of moving the RORB motif within the strong enhancer (top) and predicted contribution of the RORB motif to activity (bottom). Horizontal dashed lines represent the activity (top) and motif contribution (bottom) of the wild-type sequence. **f**, Effect on activity of swapping the strong enhancer RORB motif into the inactive sequence at different positions. Error bars show standard deviation of one sequence across 3 replicates. C, CRX motif; R, RORB motif.

To identify causal differences between these two sequences, we used the regression CNN to predict the importance of each nucleotide to the activities of both sequences (**Fig. 2a**). Strikingly, only the NRL site in the strong enhancer was scored as important, despite being identical to the NRL motif in the inactive sequence. The importance assigned by the model to the surrounding context of the NRL motifs also differed. In the strong enhancer, the NRL motif was flanked by high-importance nucleotides that form a near-optimal match to a Nuclear Receptor (NR)1-family motif (**fig. S6**). NR1-family TFs include factors that are known to interact with CRX, NRL, and other cell type-specific TFs in photoreceptors (*59, 85*), but this motif is recognized by a clade of nuclear receptor DNA-binding domains that is distinct from that of more well-characterized, photoreceptor-specific nuclear receptor TFs. The NR1-family motif was not detected as globally enriched among CRX-bound strong enhancers by traditional motif-enrichment analyses (*20*).

To test the model predictions, we performed an MPRA-based perturbation analysis (**fig. S7a**). The activity of the strong enhancer decreased when the CRX or NRL motifs were individually scrambled, and scrambling both sites abolished nearly all activity. Thus, both the CRX and NRL sites contribute to the activity of the strong enhancer, as predicted by the model. Scrambling the NR1-family motif led to a near-total loss of activity, validating the model prediction that this site is important. As predicted by the model, other regions of the enhancer had little or no effect on activity when scrambled. We next tested whether the NR1-family motif in the strong enhancer was sufficient to confer activity on the inactive sequence. We swapped the sub-sequence containing the NR1-motif into the inactive sequence, producing a chimeric sequence that was nearly as active as the original strong enhancer (**Fig. 2c**). Other chimeric sequences that did not include the region with the NR1-family motif did not lead to activation. Moving the inactive NRL motif to the same position as the NRL motif in the strong enhancer was not sufficient to confer activity on the inactive sequence, confirming that NRL positioning alone is not the causal difference between the active and inactive sequences. The CNN predicted that the NR1-family site in the chimeric sequence had an independent effect as well as a synergistic effect with the CRX site, because the importance assigned to the CRX and NRL sites in the inactive sequence both increased in the chimeric sequence (**fig. S7b**). These results demonstrate that the model discovers functionally relevant motifs, even when those motifs fail to reach statistical significance in a global enrichment analysis.

The above results suggested that the contribution of the NRL motif is affected by its proximity to the NR1-family motif. We analyzed this phenomenon by using the model to predict the importance of both NRL and NR1-family motifs as the NRL motif position was moved (**Fig. 2b**, bottom). The model predicted that the importance of the NRL motif largely decreased when moved away from its natural position, while the importance of the sequence positions containing the NR1-family motif remained high. An exception to this trend was predicted when moving the NRL motif into a position that disrupted the NR1-family motif, resulting in a loss of importance at this sequence position. Placing the NRL motif at position 128 created a cryptic NR1:NRL dimer motif (**fig. S7c**) which was predicted by the model to restore activity. We experimentally tested the predictions of the model and confirmed that sliding the NRL motif into the position of the NR1-family motif decreases CRE activity, while creating the cryptic dimer motif at position 128 restores activity (**Fig. 2b**, top). These results suggest a regulatory grammar that governs the activities of the NR1-family and NRL motifs. In the strong enhancer, the NRL motif is decommissioned if it is moved out of context, but the enhancer can tolerate this loss because of the independent contribution of the NR1-family motif. But if the NRL site moves to a location that disrupts the NR1-family motif, then the enhancer can no longer function. We conclude that the strong enhancer is composed of both a modular (NR1) and a context-dependent (NRL) motif. So long as these constraints are satisfied, the same activity can be achieved with multiple combinations and arrangements of sites, providing a flexible grammar for constructing and editing an enhancer.

We tested a second active/inactive sequence pair, each containing a central CRX motif and a motif for the nuclear receptor RORB (**Fig. 2d**). In the inactive sequence, the model predicted that neither CRX nor RORB motif was important. Surprisingly, in the strong enhancer, the model predicted that the CRX motif was not important, but the RORB motif was. These predictions were confirmed by experimental motif perturbations (**fig. S7d**). Tests of chimeric sequences showed that swapping the strong enhancer RORB motif into the inactive sequence was sufficient to confer high activity only if the RORB motif was placed 3’ of the CRX motif (**Fig. 2f**). When simulating placing the RORB motif in different positions of the strong enhancer, the model predicted that the RORB motif would remain active when positioned 3’ of the CRX motif, but that it would lose activity when positioned 5’ of the CRX motif. Experiments confirm this prediction (**Fig. 2e**). These results show that the model correctly identified a general positional requirement of the RORB motif in this strong enhancer. Taken together, these analyses of active/inactive sequence pairs confirm that the model learned critical features of sequence context that distinguish functionally non-equivalent motifs.

### The model learned relative affinities and known contextual features affecting the activity CRX motifs

The regression CNN learned previously known properties of CRX binding sites, validating the representation of the *cis*-regulatory grammar learned by the model. First, the model correctly learned the relative affinities of different CRX motif sequences. We compared motif-level importance scores assigned by the model to the measured effects of individually perturbing CRX motifs in 68 CRX-bound genomic enhancers (*82*). The model-assigned importance scores correlate with CRX motif affinity (TAATCC > TAATCD > TAAG > TGAT, where D = G/A/T, **Fig. 3a**, left panel). The relative order of the predicted importance scores for different motif variants matched the observed changes in CRE activity (**Fig. 3a**, right panel). Notably, the motif sequence TGAT is a potential CRX binding site with very low affinity (*86*), and was frequently assigned negative importance by the model. This motif variant often showed a repressive effect in the MPRA, suggesting that TGAT sites may be bound by a repressor, rather than CRX, an effect that was captured by the model.

**Figure 3:**
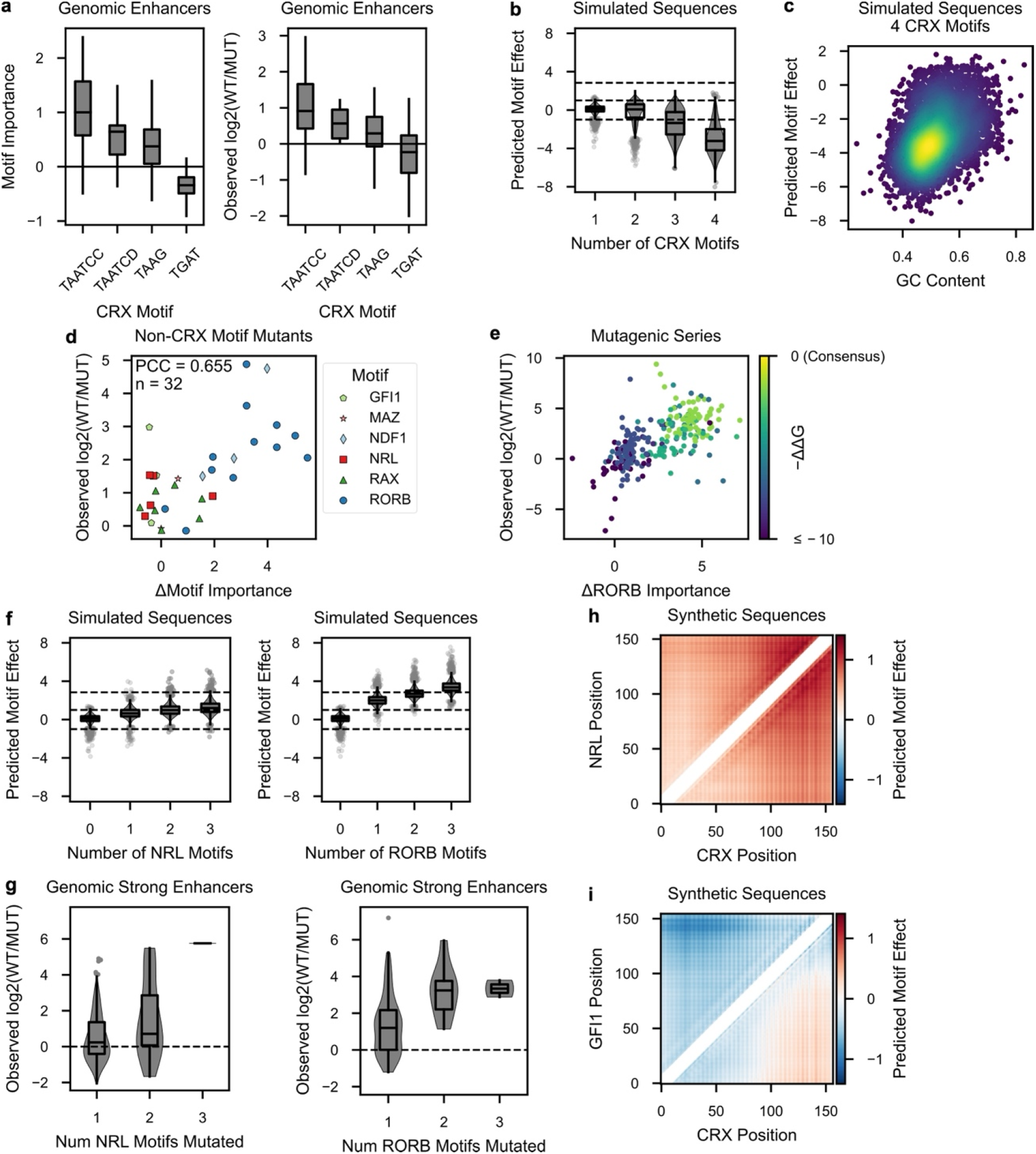
Regression CNN reveals a global photoreceptor *cis*-regulatory grammar. **a**, Predicted CRX motif importance by motif sequence (left), and change in activity when CRX motifs are individually mutated (right) in 68 genomic enhancers. “D” represents a G, A, or T at that position. **b**, Predicted effect on regulatory activity of increasing numbers of CRX motifs in simulated sequences. Horizontal dashed lines denote activity class boundaries. **c**, Directional effect of 4 CRX motifs is correlated with background GC content. Each dot represents a different background sequence. Warmer colors denote higher point density. **d**, Change in predicted motif importance versus observed effect when non-CRX motifs are individually mutated in the wild-type enhancers of the mutagenic series test set. **e**, Change in predicted RORB motif importance versus observed effect when motifs are mutated in the mutagenic series test set. Predicted motif importance corresponds with motif affinity, indicated by the color map representing the motif affinity relative to the consensus binding sequence. **f**, Predicted effect of adding NRL (left) or RORB (right) motifs to simulated sequences containing one CRX motif. Horizontal dashed lines denote activity class boundaries. Zero on the x-axis denotes the effect of one CRX motif. **g**, Effects of scrambling all NRL (left, n = 219) or RORB (right, n = 189) motifs in a set of genomic strong enhancers (*20*), stratified by the number of motifs scrambled. Dashed line indicated basal promoter activity. **h-i**, Predicted effects of a CRX motif and an NRL **(h)** or GFI1 **(i)** motif at all possible positions. Each pixel corresponds to the mean predicted effect of the two motifs in >4,600 different background sequences. White diagonal denotes excluded arrangements where the motifs would have overlapped. The basal promoter is at position 164. Colorbar is the same for both heatmaps.

We next examined whether the model learned the known repressive effect of homotypic clusters of CRX sites (*16, 20, 87*). Using an *in silico* perturbation analysis (*88*), we quantified the predicted effect of increasing the number of CRX motifs from one to four in a set of 4,658 randomly generated background elements (**Fig. 3b** and **fig. S8a**). The contribution of the CRX motifs was calculated by comparing the difference in predicted activities of the background sequences versus the motif-containing sequences. The model predicted the known repressive effects of multiple CRX sites previously observed in both genomic and synthetic CREs (*16, 87*). MPRA experiments with synthetic CREs show that sequences with four copies of the CRX motif are largely repressive (*16*), in agreement with the model predictions.

Finally, the model captured the positive influence of high GC nucleotide sequence context on the activities of CRX sites. The importance assigned by the model to four CRX sites in the *in silico* analysis increased as the surrounding GC content increased (PCC=0.387), even though GC content itself is not predicted to independently influence activity (**Fig. 3c** and **fig. S8b**). Among genomic CRX ChIP-seq peaks with varying numbers of CRX sites, the Pearson correlation between measured activity and GC content is 0.23 (*13*). The above results show that the model learned known contextual features that affect the function of CRX motifs.

### The model learned the role of other TFs enriched in photoreceptor CREs

Sequences in the training dataset all contain CRX motifs, but most sequences also contain motifs for other cell type-specific TFs that interact with CRX. We examined whether the model learned to distinguish functionally non-equivalent instances of motifs for these additional TFs. The 29 strong enhancers of the mutagenic series test set were selected because they contain motifs for six lineage-specific TFs enriched in CRX-bound CREs (*20*). In the mutagenic series, motifs for these TFs were mutated individually, allowing us to test model predictions about the importance of individual occurrences of these motifs. We compared the model-assigned importance scores for each non-CRX motif against the measured effect of perturbing that motif. Motif-level importance scores were highly correlated with the measured effects for motifs for these six additional TFs (PCC = 0.655, **Fig. 3d**). RORB sites in particular were assigned a wide range of importance scores by the model, and they exhibited a corresponding range of effect sizes when mutated. To further examine the predicted effects of RORB motifs, we increased our sample size by considering not only the single mutations of the 29 wild-type strong enhancers in the mutagenic series, but also RORB motif mutants made within the context of mutations to other motifs in these sequences (n = 258). We found that the model assigned importance scores to different RORB motif variants that correspond with their relative affinities. The model’s predictions of the effect of mutating different versions of RORB motifs were correlated with the observed effects (PCC = 0.619), with high-affinity sites showing the largest effects and low-affinity sites showing the weakest effects (**Fig. 3e**).

The model learned the relative effect sizes of NRL and RORB motifs on the activity of sequences containing a CRX motif. An *in silico* perturbation analysis of background sequences that contain one central CRX motif predicted that adding RORB motifs to a sequence has a stronger positive effect on *cis*-regulatory activity than the addition of NRL motifs (**Fig. 3f** and **fig. S8c**). These predictions are consistent with experimental data showing that the loss of RORB motifs generally causes a greater drop in activity than the loss of NRL motifs (**Fig. 3g**). The model also correctly predicted the repressive effects of GFI1 sites (**fig. S8d**), consistent with our previous finding that GFI1 sites are enriched in genomic CRX-bound silencers (*20*). The results above show that the model learned the relative average contributions of several different cell type-specific TF binding motifs. Explicit information about TF binding motifs was not provided as part of the model training, but was instead learned by the model by iteratively training on informative data consisting only of DNA sequence and measured activity. These results validate our active learning strategy as an effective method to generate the informative training data to learn the complex interactions between TF binding motifs that determine cell type-specific *cis*-regulatory activity.

While the global performance of the model shows that it has not yet learned the full *cis*-regulatory grammar of developing photoreceptors, the results above show that the model accomplished the key goal of learning to distinguish between functionally non-equivalent binding motifs for multiple cell type-specific TFs. This suggests that the model in its current state can serve as a hypothesis generator about different features of photoreceptor *cis*-regulatory grammar. We generated hypotheses about motif order and spacing by conducting an *in silico* perturbation analysis that involved systematically varying the positions of one CRX and one NRL motif in a set of >4,600 background sequences, producing a total of >100,000,000 unique predictions. This analysis predicts that the activity of CRX and NRL motifs increase as they are moved closer to the basal promoter (**Fig. 3h**). The model also predicts a synergistic effect between CRX and NRL at certain spacings (evident in the diagonal stripes in **Fig. 3h**). In an analysis of motifs for CRX and GFI1, the model predicts that sequences with GFI1 and CRX motifs are largely repressive (**Fig. 3i**). This repressive effect is eliminated when the CRX motif is within ∼65 bp of the basal promoter and the GFI1 motif is placed in a more distal position. However, when the GFI1 motif is moved close to the CRX motif, the sequences are predicted to be repressive, suggesting that the model infers that GFI1 acts through short-range repression (*89*). This analysis shows how the model can be used to generate hypotheses about complex, higher order interactions between TF binding sites which would be difficult to identify in the absence of a guiding model.

### Benchmarking alternative uncertainty sampling and sequence generation strategies in the active learning cycle

Our results show that an active learning strategy successfully improves model performance, and that our model of photoreceptor *cis*-regulatory grammar achieved the key goal of distinguishing functionally non-equivalent TF binding sites. We examined the effectiveness of specific features of our active learning workflow by comparing alternative strategies for generating and sampling candidate sequences. It is axiomatic that machine learning benefits from larger data sets, so we tested the specific effect of uncertainty sampling by comparing it to the effect of increasing the amount of training data through random sampling. We randomly sampled 6,335 sequences from the filtered candidate pool of Round 3, pooled them with the prior training data (Rounds 1 and 2) independently of 4,438 sequences picked by uncertainty sampling, and then retrained the SVM and CNN models. Models trained on the uncertainty sampling dataset outperformed those trained on data that included the randomly sampled sequences (**Fig. 4a**), even though the uncertainty sampling dataset was 11% smaller (Rounds 1 + 2 + uncertainty sampled data = 17,137 sequences; Rounds 1 + 2 + randomly sampled data = 19,034 sequences). We conclude that uncertainty sampling produces more informative training data than random sampling.

**Figure 4:**
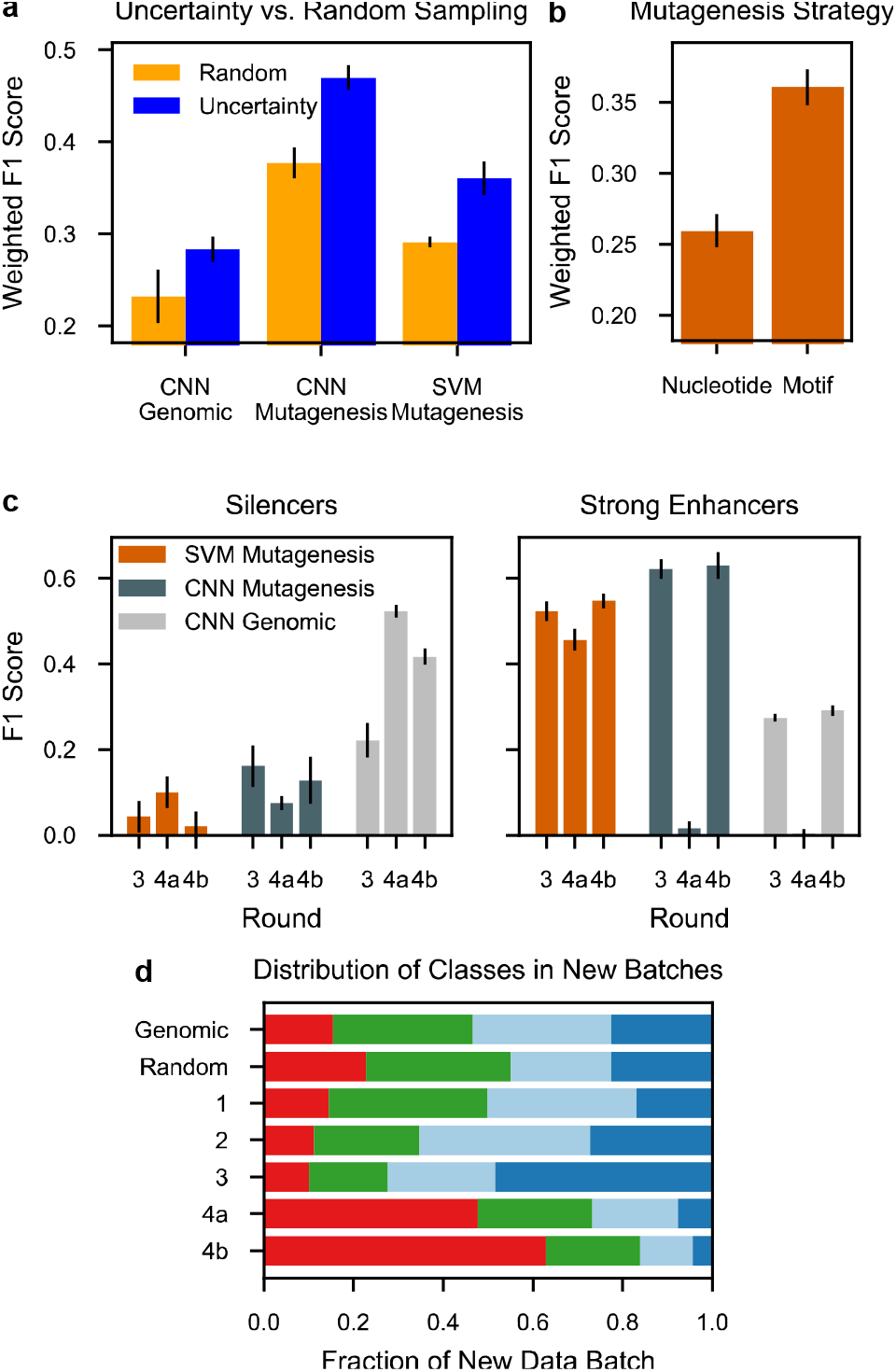
Impact of sampling and *in silico* perturbation strategies on model improvement. **a**, Performance of models when trained on new data from random or uncertainty sampling that are pooled with training data from Rounds 1 and 2. **b**, Performance of SVM predictions on the mutagenic series when trained on an equal number of sequences generated using nucleotide-based perturbations or motif-based perturbations. Training data included Round 1 sequences. **c**, Comparison of the impact of alternative sampling strategies (4a, entropy sampling; 4b, margin sampling) on model performance on silencers and strong enhancers. **d**, Variation in the distribution of activity classes in different rounds of active learning. Rows show activity classes of newly generated training data for each round (dark blue, strong enhancer; light blue, weak enhancer; green, inactive; red, silencer). All error bars are the same as in **Fig. 1c**.

The *in silico* perturbation strategy used to generate new sequences had an important effect on the utility of the training data. In Round 2, we generated candidate sequences by randomly changing 12% of the nucleotides in the sequences from the Round 1 training dataset. After completing the remaining active learning steps of Round 2, we observed no improvement in performance by the SVM (**Fig. 1c**). The CNN, which did not generalize beyond the Round 1 genomic training data, performed worse than random guessing on the mutagenic series, but achieved modest performance on the genomic test set (**Fig. 1c**). For Round 3, we changed strategies and generated candidate sequences using a motif-centric perturbation strategy in which cell type-specific motifs were added, subtracted, or moved. Models trained on the Round 3 dataset showed major improvements in global performance on the mutagenic series, while the CNN performance on the genomic test set showed little change (**Fig. 1c**). We directly compared the performance of the two sequence generation strategies by training the SVM with Round 1 and either the nucleotide-based perturbations (Round 2) or the motif-based perturbations (37% of Round 3 to equalize the dataset size), and found that motif-based perturbation led to superior performance (**Fig. 4b**). For Round 4, we used motif-centric perturbations to generate candidate sequences. Notably, both sequence generation strategies produced enough uncertain candidates for the sampling step, but these results show that not all uncertain candidates are equally informative. The success of the motif-centric perturbation strategy suggests that it is important to account for biological knowledge when designing candidates.

We next compared two methods of uncertainty sampling. In Rounds 2 and 3 we employed “entropy” sampling, which uses Shannon entropy as the uncertainty metric. Entropy for an input sequence reaches its maximum when the SVM or CNN four-way classifier assigns equal probabilities of belonging to each class. Entropy sampling thus selects candidate sequences for which the model makes no strong predictions. By Round 3, we noted that the models performed well on strong enhancers and only modestly on silencers (**Fig. 4c**). After one more round of entropy sampling (Round 4a), we observed some improvement in the classification of silencers, but the CNN overfit the data and showed a dramatic drop in performance, particularly on strong enhancers **(Fig. 1c, Fig. 4c, fig. S3**). This suggests an episode of “catastrophic forgetting” (*90*), in which learning information about silencers caused a loss of information about enhancers. We thus tested an alternative uncertainty sampling method with a parallel round of active learning (Round 4b). We performed uncertainty sampling using margin uncertainty as the metric, which is highest when the two most likely activity classes have similar probabilities. We targeted candidate sequences for which “silencer” was one of the two most probable classes. The Round 4b models avoided the performance loss on strong enhancers and the CNN improved its performance on the silencer-enriched genomic test (**Fig. 1c, Fig. 4c**). These results suggest that entropy sampling works well in early rounds of active learning when a model is relatively naive, but that margin sampling works well in later rounds when it is necessary to improve model performance on certain classes of predictions.

Our results suggest that training datasets are more effective when they include more active sequences (i.e. silencers and strong enhancers) and fewer inactive sequences. At each round, the set of untested candidate sequences picked by uncertainty sampling turned out to be enriched for strong enhancers or silencers when assayed, and contained fewer inactive sequences than the genomic dataset (**Fig. 4d**, compare Rounds 2-4b with Round 1). In contrast to the dataset picked by uncertainty sampling, the set of sequences selected by random sampling included more weak enhancers and inactive sequences (54% weak enhancer and inactive sequences by random sampling versus 42% weak enhancer and inactive sequences by uncertainty sampling at Round 3). Intriguingly, the enrichment of active classes changed at different rounds of active learning (**Fig. 4d**). Approximately half of the new training data for Round 3 consisted of sequences that, when assayed, were strong enhancers. In Round 4 more than half of the new training examples were silencers when assayed, possibly because key features of strong enhancers had been learned during earlier training rounds (**Fig. 4c**). These results suggest that training datasets enriched for active rather than inactive sequences are more informative, and that an important advantage of active learning is that it samples candidate sequences that are more likely to be active, compared to both randomly sampled and genomic datasets.

### Active learning outperforms random sampling on a human cell culture dataset

We tested active learning in a second cellular system by training a CNN on a published genome-wide MPRA dataset for enhancer activity in K562 cells (*91*). Because this dataset does not include silencer activity, we trained a binary classifier to distinguish the most active enhancers (top 20%) from inactive sequences (bottom 50%). We began by testing whether the effectiveness of uncertainty sampling varied when the size of the initial training dataset and the batch size of the sampling round were systematically varied. Using 10-fold cross-validation, we assigned each chromosome to the training, validation, or test set. From the ∼46,000 training sequences, we randomly assigned 3,000, 4,000, or 5,000 as initial training data and treated the remainder as “unlabeled” data **(Fig. 5a)**. After training an initial CNN binary classifier, we performed a single round of either entropy or random sampling by picking 1,000, 3,000, or 5,000 “unlabeled” sequences that we added to the training data with their known label, and then we retrained the CNN. Entropy sampling consistently improved the CNN and outperformed random sampling regardless of the batch size or the amount of initial training data **(Fig. 5b** and **fig. S9a)**, demonstrating that active learning is effective for training models in multiple cellular contexts.

**Figure 5:**
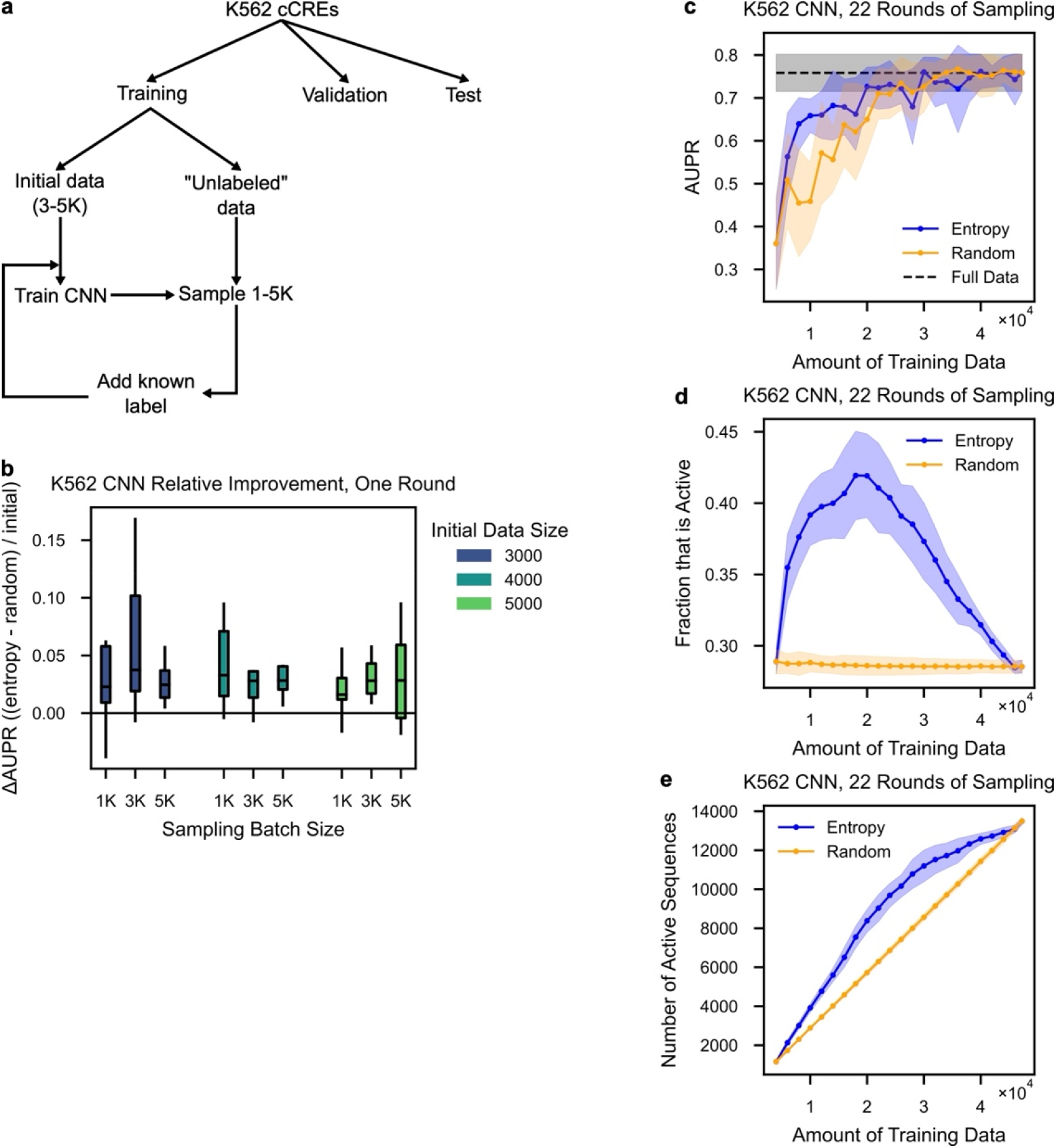
Application of active learning to K562 cells. **a**, Schematic of benchmarking experiments using K562 candidate CREs (cCREs). At each round, the validation chromosomes are used to monitor CNN training and the test chromosomes are used to evaluate final performance. **b**, Relative improvement of CNN for entropy sampling vs. random sampling compared to the initial performance, as measured by the area under the precision recall curve (AUPR) on held-out test chromosomes. Each box represents 10-fold cross-validation for a set of starting conditions. **c-e**, Comparison of multiple rounds of entropy vs. random sampling for **(c)** CNN performance, **(d)** fraction of cumulative training data that are active sequences, and **(e)** total number of active sequences sampled. Lines denote the mean from 10-fold cross-validation, shaded areas denote one standard deviation, and black denotes CNN performance with the full dataset.

We next asked whether training with active learning could achieve model performance approaching the performance of a model trained on the full dataset, but with less data. We began by training a binary classifier on an initial dataset of 4,000 sequences, then sampled batches of 2,000 at a time until all available data was exhausted (**Fig. 5c**). Entropy sampling was more efficient and outperformed random sampling until model performance approached that of the model trained on the full dataset. Model performance doubled within five rounds, and performance approached the upper bound after using only ∼40% of the full training dataset (**Fig. 5c** and **fig. S9b**). As was observed with active learning in the mouse retina, the “unlabeled” training examples picked by entropy sampling from the K562 data were strongly enriched in high-activity sequences, until those sequences were exhausted from the set of sequences available to sample (**Fig. 5d, e**). The performance of the model improved the most in rounds where entropy sampling produced a dataset enriched for high-activity sequences. In the later rounds, when there were fewer high-activity sequences remaining in the “unlabeled” pool, the gains in model performance were attenuated and performance no longer improved monotonically from one active learning round to the next. These results confirm that active learning is broadly effective in different cellular systems, and that a dataset enriched for active sequences is more informative for model training. In these benchmarking experiments with an existing dataset, the pool of sequences that could be sampled was limited, and eventually high-activity sequences were exhausted. However, in real-world scenarios, the training dataset can be indefinitely increased using synthetic DNA sequences and functional genomics technologies such as MPRAs. Thus, informative, active sequences can be added to the training data until the desired level of model performance is achieved.

## DISCUSSION

We have presented an active machine learning framework to learn the complex interactions between TF binding sites that determine the *cis*-regulatory activity of DNA sequences. We showed that active learning more than doubled the performance of models trained on a single round of genomic data alone. After four rounds of MPRAs in the developing mouse retina, we arrived at a model that makes accurate predictions, recapitulates prior knowledge, and reveals novel sequence features necessary and sufficient for enhancer activity in differentiating photoreceptors. Critically, the model distinguishes between functionally non-equivalent binding sites for multiple cell type-specific TFs, thus achieving the major goal of this study. Our work demonstrates how active learning effectively leverages the capacities of DNA synthesis and functional genomic assays to generate successive rounds of informative training data. This overcomes a critical limitation of the existing genomic datasets used to train machine learning models, which is that the number of natural training examples is often too small to learn the complex, higher-order interactions that cause binding sites for the same TF to have non-equivalent functions in different contexts. Our results suggest that active learning will be a potent approach for unraveling the effects of context on TF binding sites that determine their roles in cell type-specific gene expression. Given the continually decreasing cost of DNA synthesis and the ever-growing capacities of functional assays, active machine learning has broad potential across a range of applications, including large-scale perturbation studies of *cis*-regulatory grammars and model-guided design of synthetic regulatory DNA elements.

Our results show that smaller datasets generated by active learning produce the same model performance as models trained on larger, randomly sampled datasets. Active learning may be a more efficient approach because it generates training data that are enriched for positive examples of highly active sequences, whereas genomic sequences and random sequences contain a larger fraction of negative examples of inactive sequences (**Fig. 4d** and **5e**). We found that uncertainty sampling identifies unlabeled candidate sequences which, upon measurement, are more likely to fall into the strongest activity classes (strong enhancers and silencers). This suggests that the candidate sequences that are predicted by the model with the least confidence contain functionally-relevant patterns of sequence elements, and as a result these sequences are more likely to be active when measured. Such sequences may contrast with low-information inactive sequences which are easily learned by the model, and thus not among the candidates picked by uncertainty sampling. If this is true, then the complex, higher-order interactions between TF binding sites that define *cis*-regulatory grammars will not be learned from training data with a large fraction of inactive genomic or random sequences. In fact the problem of limited genomic training examples becomes even more acute if training data with a high fraction of active sequences are necessary to fully learn *cis*-regulatory grammars, because only a minority of candidate CREs are active when tested by functional assays (*13, 17, 77, 79*). Complementary approaches to training machine learning models of *cis*-regulatory grammars rely on hundreds of millions of random sequences (*35, 38*) or hundreds of thousands of genomic sequences (*34, 91, 92*) measured in a single, large-scale screen. Our work suggests that many of the sequences in these datasets may be low-information training examples, and that iteratively training models on smaller, but more informative training data will be more effective.

The most dramatic examples of functionally non-equivalent TF binding sites are sites in enhancers and silencers bound by the same TF (*16, 17, 93*–*96*). Loss of the CRX binding motif in enhancers causes a decrease in CRE activity, while loss of the same motif in silencers causes an increase in activity. Models that correctly predict the direction of effect when different instances of a TF binding motif are abolished have learned the local contextual patterns that cause these motifs to be functionally non-equivalent. However, current deep learning models of gene expression often fail to predict the direction of effect of non-coding variants (*97*–*99*). This may in part be due to a training data bias against transcriptional silencing, since MPRA experiments are rarely designed to capture silencer effects. An advantage of our MPRA in mouse retinal explants is that we measure both enhancer and silencer activity, thereby generating training data that allows the model to learn the contextual features distinguishing binding sites in enhancers and silencers. Non-coding genetic variants often change activity in a direction that is not expected (*100*), highlighting the importance of creating training data that can disambiguate between positive and negative effects of the same motif.

Uncertainty sampling is the crucial step of active learning. We implemented active learning in the context of discrete classifications (silencer, inactive, weak enhancer, and strong enhancer). Using these discrete classifications provided a straightforward way to implement uncertainty sampling using concepts from information theory. However, models that make quantitative predictions of *cis*-regulatory activity are often more useful, and it will thus be important to implement active learning in a regression setting by taking advantage of sampling strategies that rely on Gaussian and neural processes (*49, 101*–*103*). Our work also highlights the risks of relying too heavily on one sampling technique. In the initial rounds of active learning, we used entropy sampling, which generated enhancer-biased datasets that led to improved predictions of enhancers (**Fig. 4c** and **d**). In Round 4, both entropy sampling and margin sampling produced training datasets biased toward silencers, but entropy sampling resulted in catastrophic forgetting of strong enhancers (*90*), while margin sampling resulted in continued improvements to the model. Future work can protect against catastrophic forgetting by employing diversity criteria (*104, 105*) and using multiple sampling techniques in each round, including ensemble-based sampling (*106, 107*). Every round should also include some perturbations from random sampling to test whether uncertainty sampling is working as expected. The fact that motif-based perturbations outperformed random mutagenesis highlights the benefit of accounting for prior knowledge when generating perturbations, but such an approach induces a bias against discovering new sequence features. Recent advances in generative modeling (*29*), gradient-based design (*108, 109*), transfer learning (*110*), and evolutionary-inspired data augmentation (*111*) could provide a complementary approach to generating training data. Active learning is a broadly applicable and effective strategy for learning *cis*-regulatory grammars that can take advantage of these ongoing developments in deep learning.

## METHODS

### Library design

All MPRA libraries were created using oligonucleotide (oligo) synthesis from Agilent Technologies™ as previously described (*20, 81*). Every MPRA library contained 164 bp test sequences marked with unique 9 bp barcodes following the scheme: 5’ priming sequence, EcoRI, library sequence, SpeI, filler sequence, SphI, CRE barcode (cBC), NotI, 3’ priming sequence (**fig. S10**). The filler sequence is used to ensure all oligos are 230 nt for synthesis and is subsequently eliminated during cloning. In addition to this common design scheme, all libraries contained several groups of constant sequences: (1) a construct for the basal promoter alone, which is present with multiple redundant barcodes, (2) a set of 150 scrambled genomic sequences (3) 20 genomic sequences that span the full dynamic range of the assay. The MPRA libraries are described in **Supplementary Tables 1-3**.

### Library cloning

We created MPRA libraries from oligo pools using two-step cloning as described (*20, 81*) with the following modifications. Oligos were amplified through multiple PCR reactions (New England Biolabs [NEB] Q5 High-Fidelity 2X Master Mix, cat. #M0515, see **Supplementary Table 4** for primer sequences), purified from an agarose gel, digested with EcoRI-HF and NotI-HF (NEB), and then cloned into the EagI and EcoRI sites of pJK03 (AddGene #173,490) in multiple ligation reactions (NEB T4 ligase). Ligation products were transformed into either 5-alpha or 10-beta electrocompetent cells (NEB) and grown in liquid LB-Amp cultures. Plasmid pools were digested with SphI-HF and SpeI-HF and treated with Antarctic phosphatase or Quick CIP (NEB), then ligated to reporter gene inserts in multiple reactions (NEB T4 ligase). Ligation products were transformed into electrocompetent cells and grown in liquid culture from which we prepared plasmid DNA.

We next cloned the *Rho* basal promoter into the plasmid library in between the test sequence and its cognate barcode. Basal promoter inserts were prepared by amplifying the *Rho* basal promoter and *DsRed* from the plasmid pJK01 (AddGene #173,489) using the forward primer MO566 and reverse primers that add 9bp multiplexing barcodes (mBC, **Supplementary Tables 1 and 2**), purified from an agarose gel, and digested with NheI-HF and SphI-HF (NEB). Adding mBCs to the reporter gene allows us to test larger libraries by amplifying sublibraries with different primer sets and cloning each sublibrary in parallel with a unique mBC. When necessary, we could then mix sublibraries (**Supplementary Table 1**) for parallel analyses. Barcode complexity was always verified by sequencing the final library on the Illumina MiniSeq™ platform.

### MPRA in mouse retinal explants

Animal procedures were performed in accordance with a Washington University in St. Louis Institutional Animal Care and Use Committee approved vertebrate animals protocol. CD-1 IGS mice were obtained from Charles River Laboratory. Retinas from newborn (P0) mice were dissected and electroporated as described (*69*). The sex of the mice could not be determined at the P0 stage. Retinas were dissected in serum-free medium (SFM; 1:1 Dulbecco’s Modified Eagle Medium (DMEM):Ham’s F12 (Gibco, 11330-032), 100□units per ml penicillin and 100□μg□ml−1 streptomycin (Gibco, 15140122), 2□mM GlutaMax (Gibco, 35050-061) and 2□μg□ml−1 insulin (Sigma, I6634) from surrounding sclera and soft tissue leaving the lens in place. Retinas were then transferred to an electroporation chamber (model BTX453 Microslide chamber, BTX Harvard Apparatus modified as described (*112*)) containing 0.5□μg□μl−1 of MPRA library. Five retinas were pooled for each biological replicate and at least three replicates were performed for each library (**Supplementary Table 1**). Five square pulses (30□V) of 50-ms duration with 950-ms intervals were applied using a pulse generator (model ECM 830, BTX Harvard Apparatus). Electroporated retinas were removed from the electroporation chamber and allowed to recover in SFM for several minutes before being transferred to the same medium supplemented with 5% fetal calf serum (Gibco, 26140-079). The retinas were then placed (lens side down) on polycarbonate filters (Whatman, 0.2□μm pore size 110,606) and cultured at 37□°C in SFM supplemented with 5% fetal calf serum. After eight days of culture, we harvested the retinas in TRIzol, homogenized tissues with a sterile needle, and extracted RNA following the manufacturer’s protocol. RNA was treated with TURBO DNase and then reverse transcribed with SuperScript IV First Strand Synthesis following the manufacturer’s protocol. Barcodes were amplified from cDNA and plasmid pools with Q5 using primers BC_CRX_Nested_F and BC_CRX_R for 25 cycles. We performed 2 PCR reactions per cDNA sample and 1-2 reactions per plasmid pool. PCRs from the same sample were then pooled and purified. Custom sequencing adapters were added with two rounds of PCR with Q5. The final libraries were sequenced on the Illumina NextSeq or NovaSeq platform.

### Data processing

Sequencing reads were filtered for reads that contained both the cBC and the mBC (when utilized) in the correct sequence context. One sequencing library (**Supplementary Table 1**) contained a systematic error that led to N’s in positions 5, 16, 21, 27, 29, and 34. Position 5 was in a sequencing adapter, positions 16 and 21 were in constant regions, and positions 21, 27, and 34 were in the mBC region. Since the mBC could only be one of two 9 bp sequences with a Hamming distance of 8, we could still unambiguously assign mBC-cBC pairs if there are no other errors in the read outside of these 6 positions.

We removed cBCs with fewer than 50 counts in the plasmid pool or with a coefficient of variation above 0.8 across cDNA samples. Each sample was then normalized by sequencing depth; sublibraries that were cloned separately but co-electroporated were processed separately to account for any cloning batch effects. cDNA barcodes were normalized to plasmid barcode abundances; then, barcodes corresponding to the same CRE were averaged together to obtain an activity score for each CRE in each replicate. These activity scores were normalized to within-sample basal activity and then averaged together to obtain a final activity score. In cases where basal recovery was poor (median RNA/DNA barcode ratio below 0.05 in any one replicate), we calculated a pseudobasal activity using the cBCs corresponding to scrambled sequences. We took this as a reasonable approximation of basal activity because the average activity of scrambled sequences in our initial library is no different from basal. All within-library quality control summary statistics are in **Supplementary Table 5**. Activity scores were discretized into four classes using the cutoffs we previously reported (*20*).

### Filter SVM for open chromatin

We fit a *k*-mer SVM to classify rod photoreceptor ATAC-seq peaks (*60*) from the rest of the mouse genome. We used the central 300 bp of all 39,265 rod ATAC-seq peaks as positives and selected an equal number of GC-matched sequences from the mm10 genome using the script `nullseq_generate.py` from the gkmSVM package. We fit this model using LS-GKM (*113*) with parameters `-l 10 -k 6 -d 3` and five-fold cross-validation. We assessed model performance using the Area Under the Receiver Operating Characteristic (AUROC, **fig. S1**). In each round of *in silico* mutagenesis, we selected perturbations that had an LS-GKM score of 1 or greater; this score corresponds to a sequence that is beyond the maximum-margin hyperplane. Empirically, this removed at least half of all perturbations in each cycle.

### SVM classifiers of MPRA activity

For our SVM classifier of MPRA activity, we implemented a version of the *k*-mer kernel that accounts for *k*-mer position. Since all sequences in our initial training data were centered on a high-quality CRX motif, *k*-mer position effectively reflects the distance of a *k*-mer from a CRX motif. As Giguère *et al*. show (*114, 115*), the Generic String Kernel for any two strings *y,y*’is:

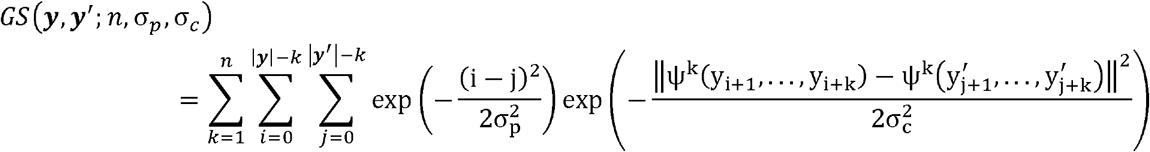

where *n* controls the maximum length of *k* -mers, σ _*p*_ controls the weight of *k* -mer position, σ _*c*_ controls the weight of *k* -mer similarity, and ψ ^*k*^ is a *k* -mer encoding function. (We use *k* wherever Giguère *et al*. used 1.)

When *σ*_*c*_ = 0, the second exp(·) becomes an indicator function that is only true if the two *k*-mers are identical. If we further remove the first summation from all 1,…,*n* possible *k*-mers and only consider *k*-mers of length n, we can rewrite the above kernel function as:

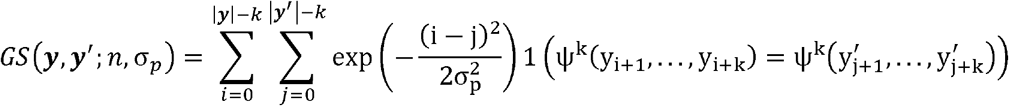

When *σ*_*p*_ =, the first exp (·)becomes constant and we recover the *k*-mer kernel (*116, 117*). We extended Giguère *et al*.’s implementation for peptide sequences to allow for DNA *k*-mers; using this implementation, we pre-computed the Gram matrix for all available data. Then, we used the `SVC` class from scikit-learn (*118*) with `probability = True` to fit our multi-class classifier. We performed a grid search over the hyperparameters *k* ∈ [6,8] and *σ*_*p*_ ∈ [0,3,10,20,50,∞] using five-fold cross-validation on our initial training data. We selected *k* = 6, *σ*_*p*_ = 10 based on the AUROC and used these hyperparameters for all future modeling.

### CNN classifiers of MPRA activity

Our CNN was designed as a multi-class classifier that uses one-hot encoded 164-bp long DNA sequence (A = [1,0,0,0], C = [0,1,0,0], G = [0,0,1,0], T = [0,0,0,1]) to predict its activity in retinal MPRAs. Our model architecture consists of two convolutional layers, a max-pooling layer, a third convolutional layer, and a second max-pooling layer; this is followed by a single fully connected layer, and finally a four-node output layer with log soft-max activation, which corresponds to the log-probability the sequence belongs to each of four classes. Every convolutional and fully connected layer is followed by batch normalization, Leaky ReLU activation, and dropout regularization. We implemented the model in Pytorch (*119*) and used Selene (*120*) to train the model with Stochastic Gradient Descent (learning rate = 0.0001, momentum = 0.9, weight decay = 10^−6^), negative log likelihood as a loss function, and a batch size of 64. We fit the model for 500 epochs using the default learning rate scheduler in Selene and kept the model with the lowest loss on a held-out validation set. We manually adjusted hyperparameters and model architectures to yield best performance on the validation set when using Round 3 as training data. We used these hyperparameters for all other datasets.

### Construction of validation set

We created a validation set for the CNN by randomly sampling 10% of our original genomic sequences and then adding all perturbations derived from those sequences in Rounds 1-3. Similarly, all perturbations derived from our test set (described below) were removed from training and validation datasets for all machine learning models. To assess model performance, we performed ten-fold cross-validation on the newly added data while holding any previous data constant. This strategy ensures that the variation in model performance is only due to variation within the new data.

### *In silico* mutagenesis for candidate perturbations

In each round, we generated a new pool of candidate sequences by perturbing sequences from the current round of training data. We defined multiple operations and performed combinations of these operations many times to generate multiple candidate sequences from each training sequence. In Round 2, the possible operations were: (1) randomly mutagenize ∼12% of the positions (the exact number of positions was chosen by sampling from a Poisson distribution), (2) insert, delete, or move a random *k*-mer, (3) create a chimera with another randomly selected sequences, and (4) randomly rearrange blocks within the sequence. We performed each of these operations multiple times. In Round 3, we implemented motif-centric perturbations based on our reference list of 8 TFs (CRX, GFI1, MAZ, MEF2D, NeuroD1, NRL, RORB, and RAX) (*20*), plus ELF1 and TBX20. We computed the predicted occupancy with μ=9 for these TFs, defined spacer sites as positions with total predicted occupancy below 0.5, and then randomly selected one of the following operations: (1) sample from one of the position weight matrices and insert that motif into a random spacer region, (2) select an occupied site and scramble it to an unoccupied state, (3) select an occupied site and replace it with a motif sampled from a different position weight matrix, (4) select an occupied site and swap it with a length-matched spacer region. We generated additional candidate sequences by systematically scrambled tiles of spacer regions. For Round 4 we excluded ELF1 and TBX20 from the motif pool due to their very low representation in the genomic sequences.

### Active learning

Our machine learning models take a 164-bp sequence *x* as input and predicts *p*(*y*_*i*_ ❘ *x*), the probability that the sequence belongs in the *i* -th activity bin, *i* = 1,…,4. Our objective is to determine which perturbations are the most uncertain to the model. Entropy uncertainty is quantified with Shannon entropy,*S*(*x*) = −. Σ *p*(*y*_*i*_ ❘ *x*) log_2_ *p* (*y*_*i*_ ❘ *x*). Margin uncertainty is defined as 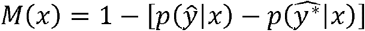, the difference in probability between the two most likely outcomes, *ŷ* and 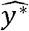. When ŷ, 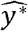 are equally likely the term in brackets is zero, so the complement represents the uncertainty.

To provide intuition for the difference between Shannon entropy and margin uncertainty, consider three cases where *ŷ*,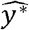 are equally likely. In the first case, a model outputs [0.25,0.25,0.25,0.25], so *S* = 2 and *M* = 1. In the second case, a model outputs [0.33,0.33,0.33,0]; once again *M* = 1, but *S* = 1.6. In the third case, the output is [0.5,0.5,0,0] and *M* is still 1, but *S* = 1. Thus, as the two most likely classes become more distinguishable from the remaining classes, the entropy can drop without a change in margin uncertainty.

In Rounds 2 and 3, we calculated probabilities with our SVM from the previous round. In Round 2, we sampled 4800 perturbations with high entropy uncertainty. In Round 3, we sampled the 13,986 perturbations with the highest entropy uncertainty. We also randomly sampled 6584 perturbations and 25 perturbations with high probability for each of the 4 classes (100 sequences total).

For Round 4, we calculated probabilities with our CNN trained in Round 3. In Round 4a, we sampled 96,190 perturbations with the highest entropy uncertainty. In Round 4b, we sampled 18,000 perturbations with entropy uncertainty below 1.8 and high margin uncertainty for silencers. Of these, 6000 were on the silencer side of the strong enhancer vs. silencer margin, 6000 were on the strong enhancer side of the strong enhancer vs. silencer margin, and 6000 were on the silencer vs. inactive margin. Very few perturbations were on the silencer vs. weak enhancer margin, so we did not sample on this margin. We chose an entropy uncertainty cutoff of 1.8 because this approximately corresponds to probabilities of [0.32,0.32,0.32,0.05], which represents cases where one outcome is unlikely but the rest are equally likely. Finally, we sampled 500 perturbations with the highest probability of being a strong enhancer, 500 high-probability weak enhancers, 500 high-probability inactive sequences, and 1500 high-probability silencers.

For all data batches, the number of sequences sampled is larger than what is listed in the main text because not all sequences were recovered upon sequencing.

### Evaluating model performance on test datasets

The genomic test set is from a parallel study (*82*) and contains 1723 CRX ChIP-seq peaks not tested in any previous study. We normalized activity scores to the basal *Rho* promoter and grouped strong and weak silencers into one silencer category, but otherwise did not re-process the data.

The mutagenic series is based on 29 CRX ChIP-seq peaks that are strong enhancers in our original Round 1 data (*20*); 17 of these become weak enhancers when all CRX motifs are mutated, while 12 remain strong enhancers. For each sequence, we computed the predicted occupancy for our reference list of 8 TFs. Then we selected all possible combinations of motifs and scrambled each motif and 3 bp of flanking sequence until its predicted occupancy was below 0.01. We tested these 727 sequences in the same library as Round 3 (**Supplementary Table 1**).

We utilized the F1 score as our evaluation metric. For any class *i*, the F1 score is the harmonic mean between the precision and recall, or equivalently, 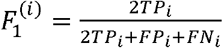 where *TP* is the number of true positives, *FP* is the number of false positives, and *FN* is the number of false negatives. The weighted F1 score is 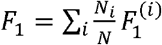 where *N* is the total number of examples and *N*_*i*_ is the number of examples belonging to class *i*.

### Additional perturbation datasets

To find inactive sequences whose motif content was similar to the strong enhancers in **Fig. 2**, we selected previously tested inactive genomic sequences with the same number and identity of motifs, and selected the sequence with the highest 6-mer similarity. We partitioned the sequences into non-overlapping blocks based on the motif positions in the inactive and strong enhancer sequences (coordinates in **Supplementary Table 6**). Then we swapped blocks individually and in combinations of either motif blocks or spacer blocks (but not both); we also moved a motif internally from its native position to its corresponding position in the other sequence before swapping blocks **(Supplementary Table 7**). Last, we moved the non-CRX motif in the strong enhancer to every other position that did not overlap with the CRX motif (**Supplementary Table 8**).

To test the effects of NRL, NeuroD1, RORB, and MAZ motifs, we scrambled all instances of these motifs in all strong enhancers in the Round 1 genomic library. Motifs were scrambled either individually or in combination with mutating all CRX motifs via point mutation. We assayed these sequences in the same library as Round 3 (**Supplementary Table 1**).

### Regression model

To perform model interpretation, we trained a new regression CNN that predicts log2 MPRA activity directly from sequence. We updated our architecture to include residual skip connections, dilated convolutions, and first-layer exponential activation (*121*). This architecture consists of three “convolution blocks,” a fully connected layer, and a final single-output node. A convolutional block consists of a convolutional layer, multiple dilated convolutional layers with residual skip connections, and then max-pooling. We trained the model at Round 4b and used the same validation set to tune hyperparameters. We trained the model using the Adam optimizer (learning rate = 0.0003, weight decay = 10^−6^), mean squared error as a loss function, and a batch size of 128. We fit the model for 50 epochs with early stopping (patience = 10, metric = Spearman) and a custom learning rate scheduler (patience = 3, decay = 0.2). Our final model was selected by training 20 initializations and selecting the one with the highest PCC on the validation set.

### Motif analysis

All motif analyses were performed using our predicted occupancy framework and μ=9 (*122*). At this value, a motif with a relative K_D_ = 3% of the consensus site has 50% probability of being occupied. To identify individual motifs, we compute the predicted occupancy of a TF and identify the positions where the predicted occupancy is at least 0.5. To identify the total number of motifs for a TF, we sum the predicted occupancy across every position of the sequence. Cartoon sequences in **Fig. 2** were generated using the predicted occupancies of our reference list of 8 TFs.

### *In silico* global importance analysis

We used Global Importance Analysis (*88*) to predict the global effect of specific sequence features on MPRA activity. We generated a background distribution by dinucleotide shuffling each of our 4658 genomic sequences and predicting their activity with our regression model. Then, we injected a fixed sequence feature in a fixed location of every sequence in our background distribution, predicted their activity with the same model, and subtracted the predictions for the background distribution. The result is the predicted log2 fold change of a sequence feature on MPRA activity, and when averaged across all sequences, represents the global importance of that feature.

### Nucleotide contribution scores

We predicted the contribution of each nucleotide to regulatory activity by calculating a sequence’s saliency map from the regression CNN followed by the Majdandzic correction method (*123*). This method has been shown to reduce noise in feature attribution maps. Motif importances scores were obtained by summing across all nucleotides that overlap predicted occupancy hits.

### Analysis of MPRA data from K562 cells

We downloaded the large-scale K562 dataset from (*91*) (Supplementary Tables 3 and 4 in their manuscript). We selected sequences that were observed in multiple replicates, had a sample coefficient of variation less than or equal to 0.75, belonged to the “putative enhancer” category, and were in the positive strand orientation. Among these sequences, we defined the top 20% as positives, the bottom 50% as negatives, and removed sequences in the 50-80th percentile.

We split the data into 10 folds based on chromosomal origin so that each fold contained approximately 10% of the data. The folds are:

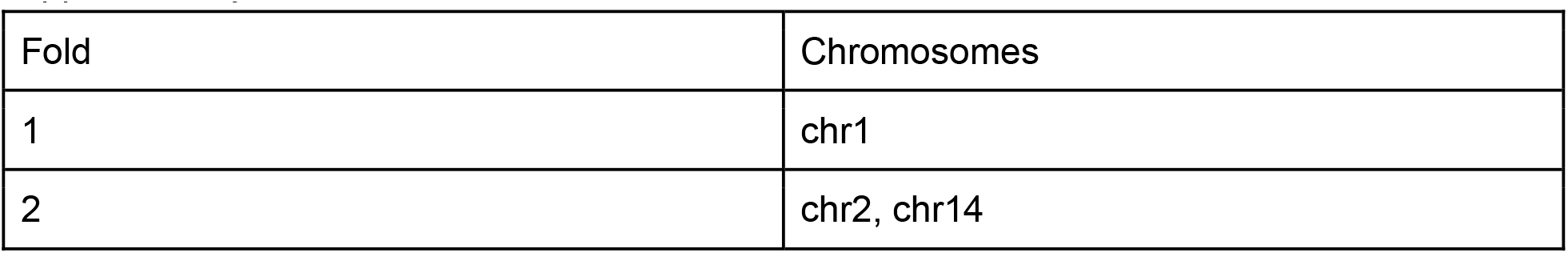
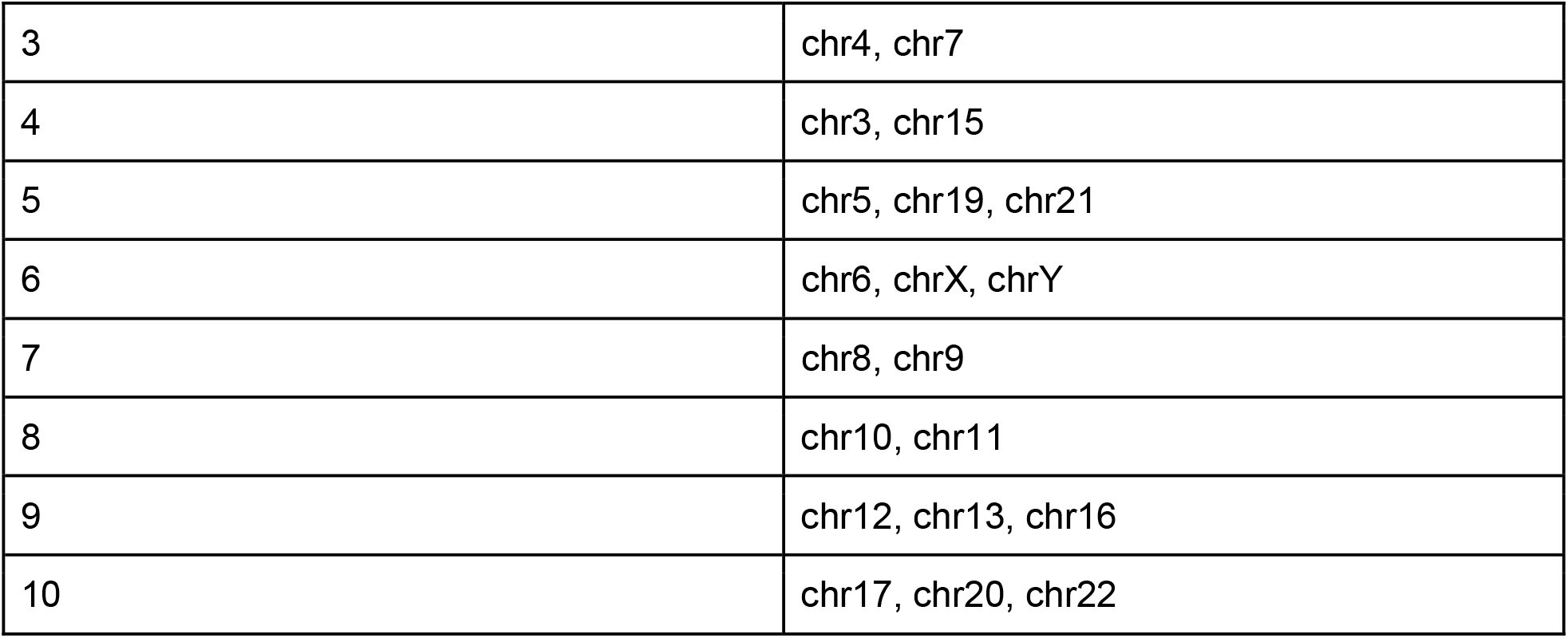

We used the same CNN architecture as our regression model, but using 230 bp input and adding a sigmoid activation function to the output to convert it into a binary classifier. We trained all models using the Adam optimizer (learning rate = 0.0001, weight decay = 10^−6^), binary cross-entropy as a loss function, a batch size of 128, and 100 epochs with early stopping (patience = 15, metric = AUPR) and the default learning rate scheduler in Selene. We kept the model with the lowest loss on the held-out validation set for evaluating on the test set and performing additional rounds of sampling.

### Statistics and data visualization

All statistical analyses and data visualization were performed in Python with Numpy (*124*), Scipy (*125*), Pandas (*126*), Matplotlib (*127*), and Logomaker (*128*). All correlations were calculated using the functions `scipy.stats.pearsonr` and `scipy.stats.spearmanr`. In all box plots, the line denotes the median, the box represents the interquartile range (25th to 75th percentile), and whiskers extend to 1.5x the interquartile range. Violin plots cover the same range as box plots, with any outliers shown as translucent dots.

### Ethics

This study was performed in strict accordance with the recommendations in the Guide for the Care and Use of Laboratory Animals of the National Institutes of Health. All of the animals were handled according to protocol A-3381-01 approved by the Institutional Animal Care and Use Committee of Washington University in St. Louis. Euthanasia of mice was performed according to the recommendations of the American Veterinary Medical Association Guidelines on Euthanasia. Appropriate measures were taken to minimize pain and discomfort to the animals during experimental procedures.

## Data availability

Sequence data that support the findings of this study have been deposited with the NCBI Gene Expression Omnibus (GEO) with the primary accession codes GSE165812 (Round 1 library) and GSE241353 (https://www.ncbi.nlm.nih.gov/geo/). Processed MPRA data files used in this study are available at [https://github.com/barakcohenlab/CRX-Active-Learning].

## Code availability

Code used to process all MPRA data, to train machine learning models, and to reproduce the results of this study are available at [https://github.com/barakcohenlab/CRX-Active-Learning]. Our implementation of the Generic String Kernel is available at [https://github.com/barakcohenlab/preimage]. Our fork of the Selene package with support for custom early stopping and learning rate decay is available at [https://github.com/rfriedman22/selene].

## ACKNOWLEDGEMENTS

We thank Roman Garnett, Peter Koo, and Kathleen Chen for machine learning and software help; members of the Cohen laboratory for helpful discussions and critical feedback on the manuscript; Jessica Hoisington-Lopez and MariaLynn Crosby in the DNA Sequencing Innovation Lab for assistance with high-throughput sequencing; and Brian Koebbe and Eric Martin for computing cluster support. This work was supported by National Institutes of Health grants R01 GM121755 to M.A.W.; R01 GM092910 to B.A.C.; R01 EY030075, HL149961, and MH122451 to J.C.C.; and F31 HG011431 to R.Z.F.

## AUTHOR CONTRIBUTIONS

R.Z.F. conceived the project. R.Z.F., M.A.W., and B.A.C. designed the overall project. R.Z.F. performed all data processing and quality control. R.Z.F. and S.L. implemented the machine learning methods. R.Z.F., A.R., S.L., Y.W., L.T., and D.L. performed computational analyses. R.Z.F. and D.G. cloned libraries and prepared samples for sequencing. C.A.M. and M.G. performed retinal dissections and electroporations. J.C.C., B.A.C., and M.A.W. supervised the project. R.Z.F., J.C.C., B.A.C., and M.A.W. prepared the manuscript with input from all authors.

## SUPPLEMENTARY FIGURES

**Fig. S1:**
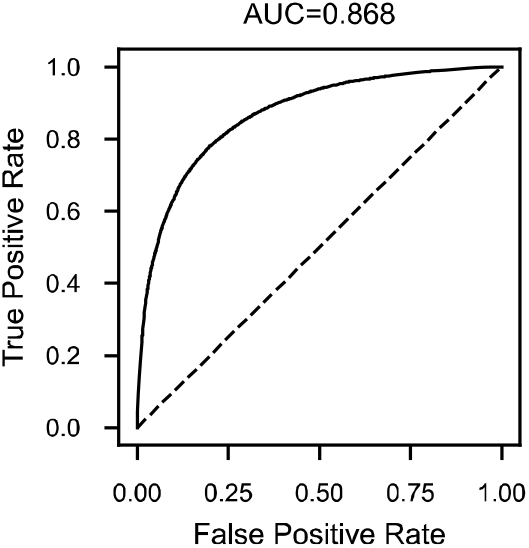
Receiver operating characteristic (ROC) curve of the filter SVM. Performance is based on five-fold cross-validation. Dashed diagonal denotes chance.

**Fig. S2:**
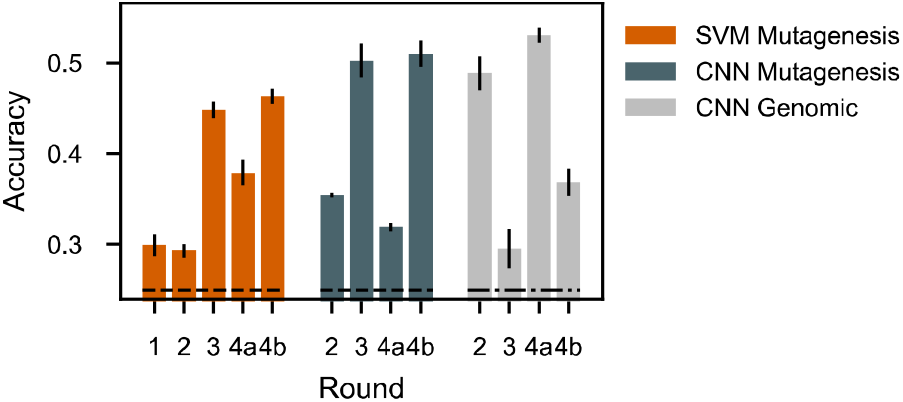
Additional performance evaluation of classifiers. Accuracy is equivalent to the micro F1 score. Horizontal dashed line represents chance for both test sets. Error bars denote one standard deviation based on ten-fold cross-validation of the newly added data.

**Fig. S3:**
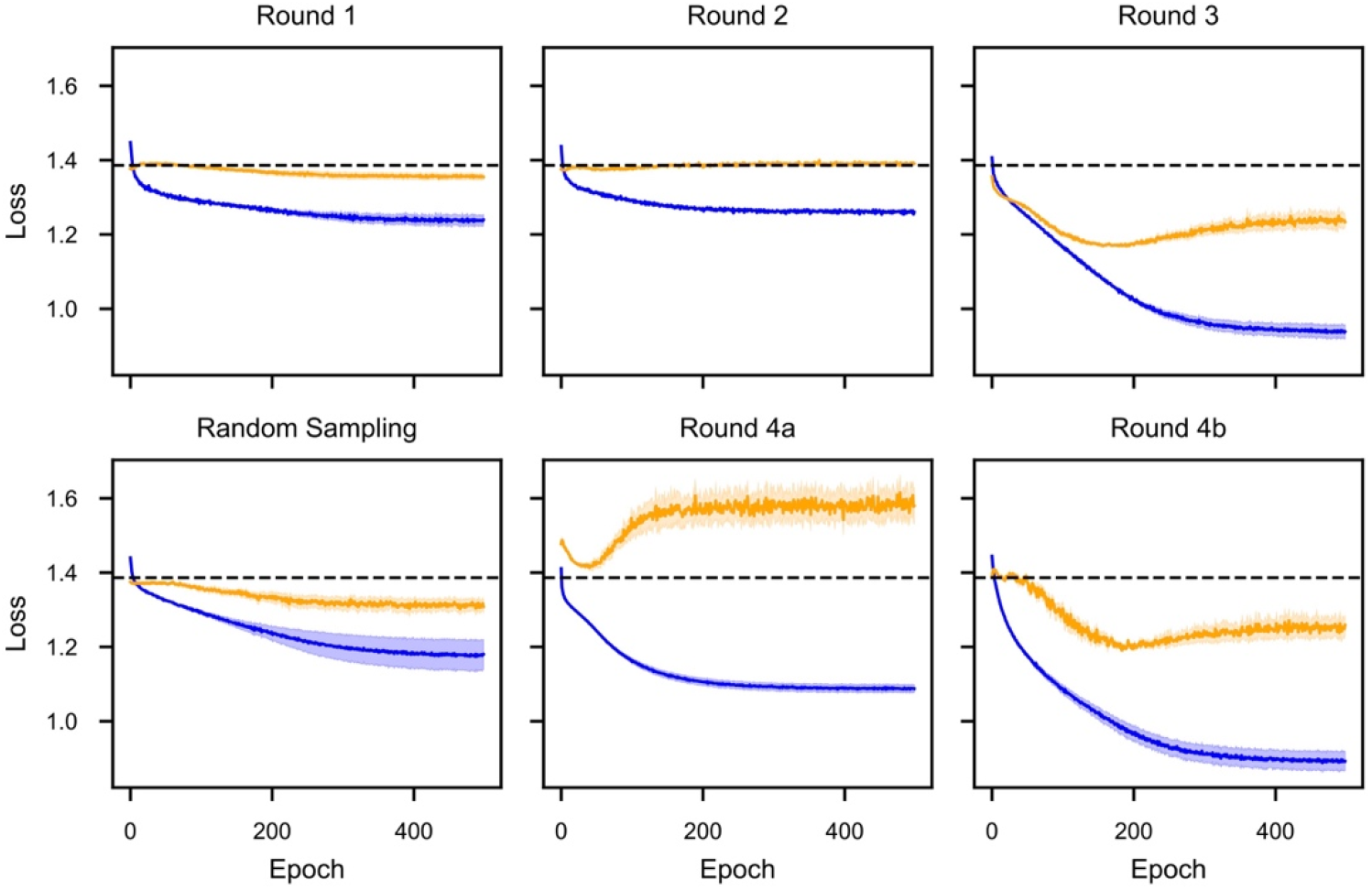
CNN classifier training history for each round of learning. Blue line denotes training data, orange line denotes validation data, horizontal dashed line denotes chance. Shaded areas denote 1 standard deviation based on ten-fold cross-validation on the newly added data. The final model parameters used for each round were selected from the epoch where the validation loss is minimized.

**Fig. S4:**
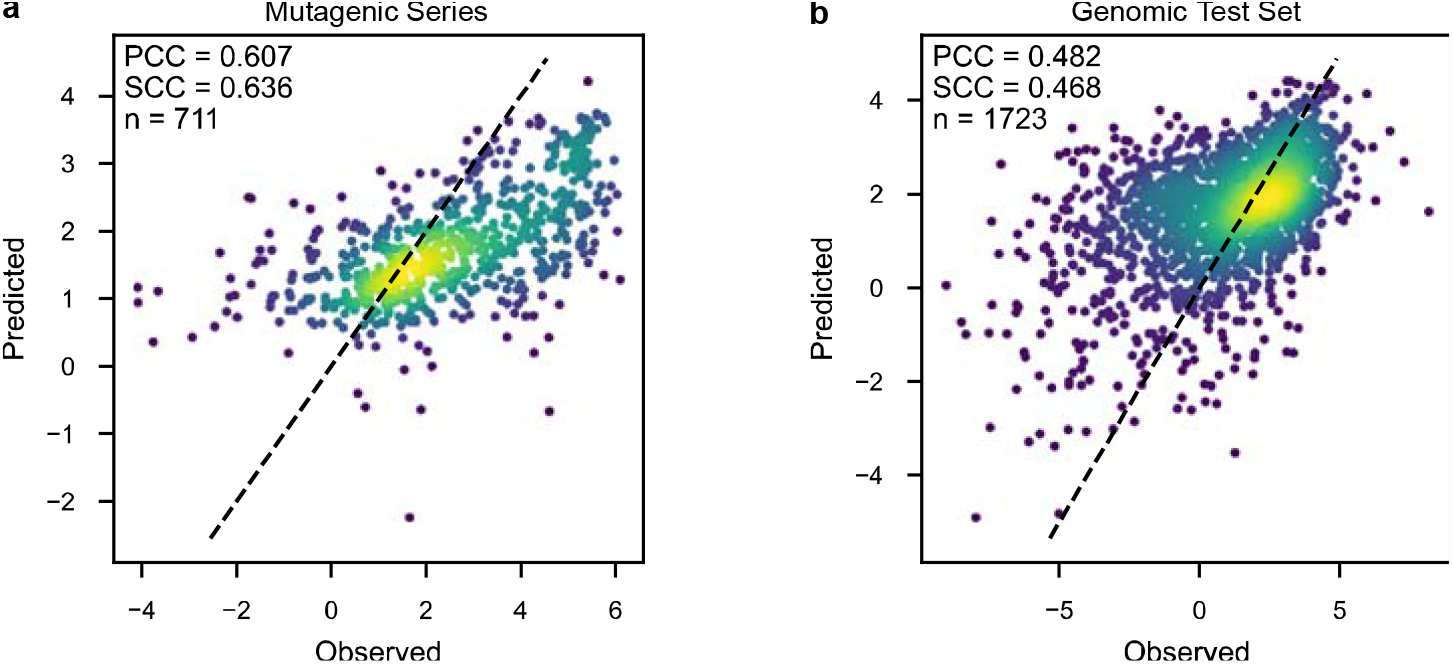
Regression CNN performance on test sets. Predicted versus observed activity from the final model (Round 4b) for the mutagenic series **(a)** and genomic test set **(b)**. Diagonal dashed line denotes equality. Cooler colors denote regions of higher density.

**Fig. S5:**
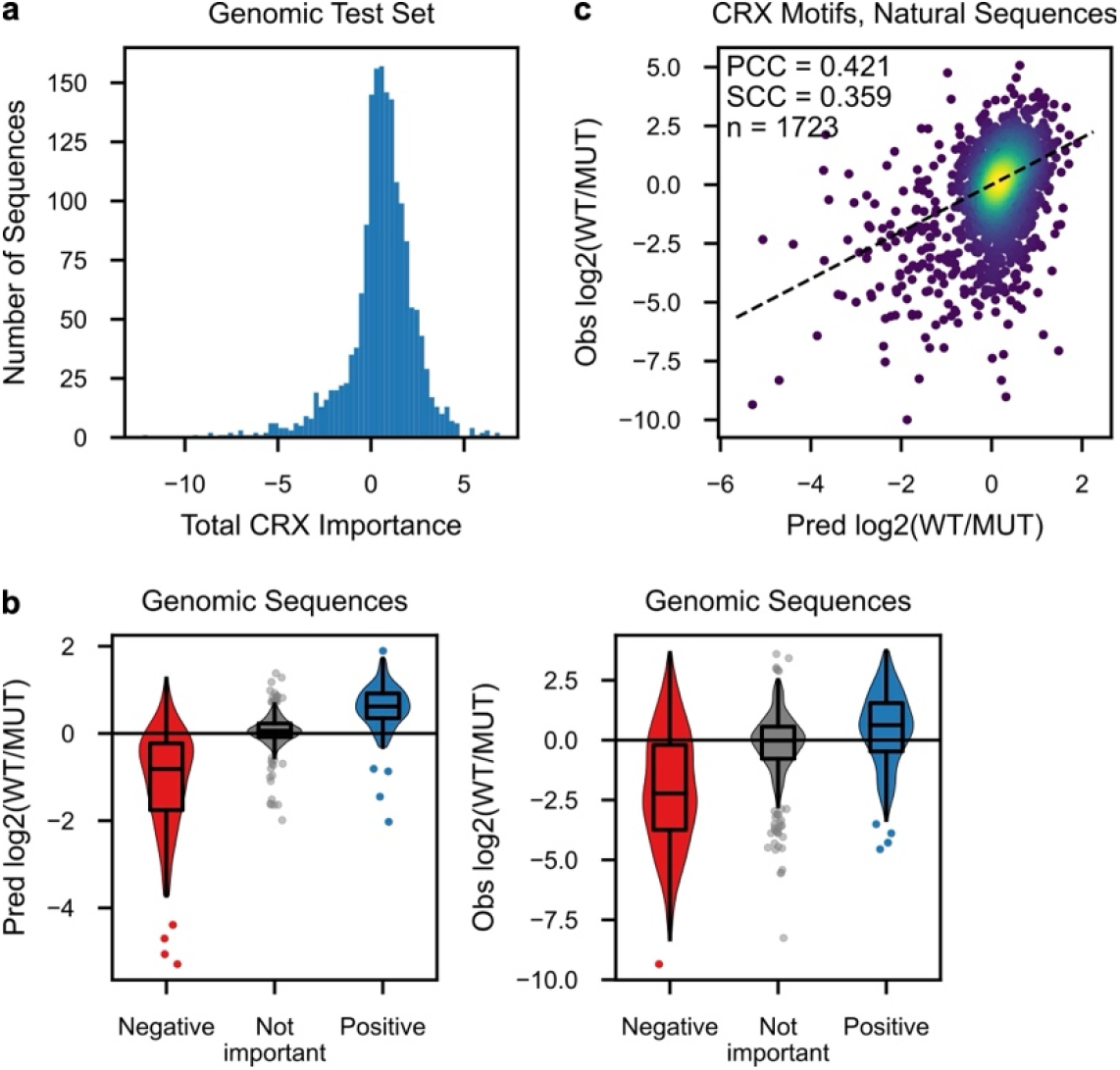
Additional validation of CRX motif importance scores. **a**, Distribution of CRX motif importance scores across the genomic test set. **b**, Predicted and observed effects of mutating CRX motifs in the genomic test set, stratified by CRX importance scores. **c**, Predicted and observed effects of mutating CRX motifs across the entire genomic test set. Diagonal dashed line denotes equality. Cooler colors denote regions of higher density.

**Fig. S6:**
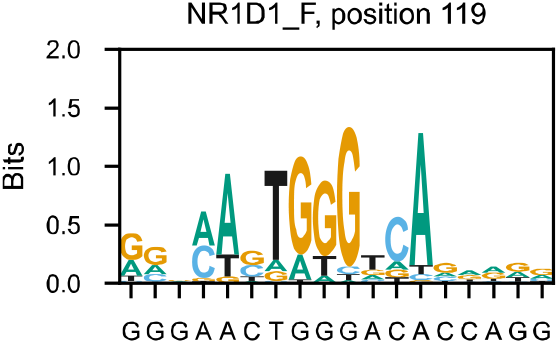
NR1-family motif match in the strong enhancer. The motif for NR1D1 is used as a representative motif of the NR1 TF family. X-tick labels indicate the sequence that matches the motif. The position and orientation of the motif is indicated in the title.

**Fig. S7:**
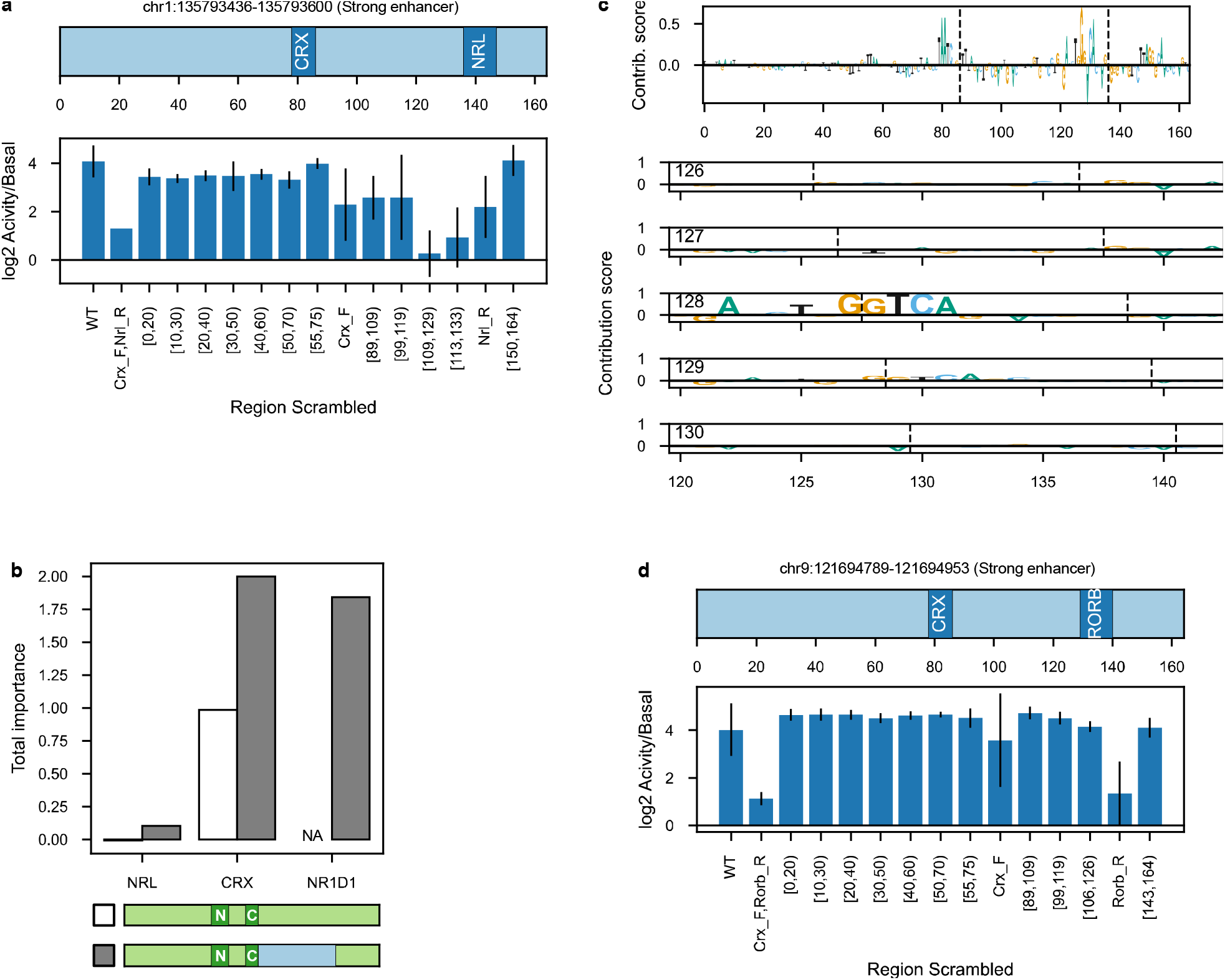
Additional validation of causal sequence differences. **a**, Observed effect of scrambling different components of the strong enhancer in **Fig. 2a-c**. Error bars show standard deviation of 3-5 independent scrambles, or 4 independent replicates of wild-type. **b**, Importance scores of the NRL, CRX, and NR1-family motif in the inactive sequence before and after swapping in region 86-136 from the strong enhancer. **c**, Relative nucleotide contribution scores when positions 86-136 from the strong enhancer (containing the NR1-family site) are swapped into the inactive sequence (top row), and nucleotide contribution scores of the strong enhancer when the NRL motif is moved to positions ranging from 126-130. The cryptic NR1:NRL site is formed when NRL is moved to position 128. Vertical dashes denote the breakpoints of the swap or the boundaries of the sliding NRL sequence. **d**, Same as **(a)** but for the strong enhancer in **Fig. 2d-f**.

**Fig. S8:**
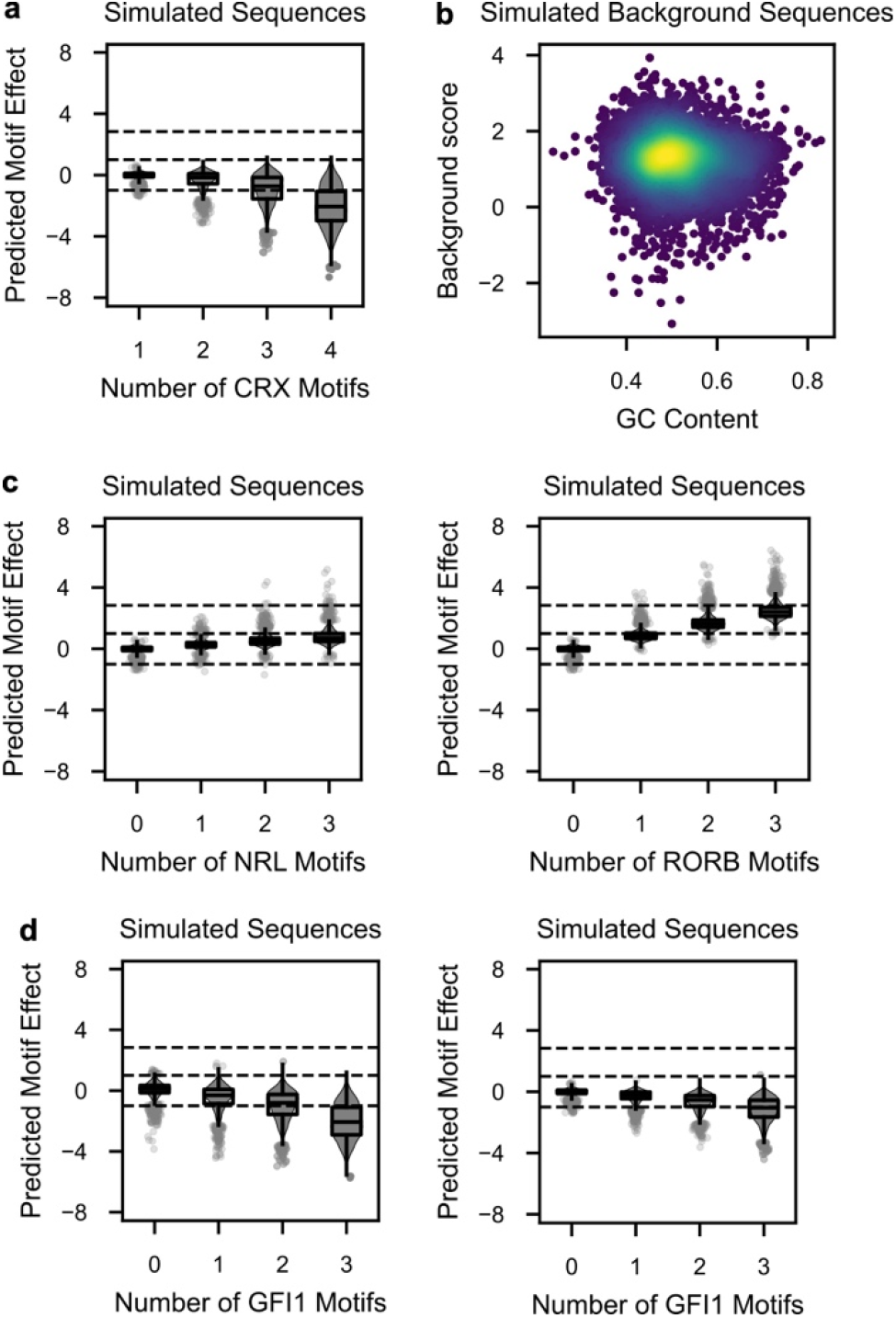
Additional *in silico* perturbation analyses. **a**, Predicted effect on regulatory activity of increasing numbers of CRX motifs in simulated sequences at a second set of positions. Horizontal dashed lines denote activity class boundaries. **b**, Lack of correlation between predicted activity of simulated background and GC content. Each dot represents a different background sequence. Warmer colors denote higher point density. **c**, Predicted effect of adding NRL (left) or RORB (right) motifs to simulated sequences containing one CRX motif at a second set of positions. Horizontal dashed lines denote activity class boundaries. Zero on the x-axis denotes the effect of one CRX motif. **d**, Predicted effect of adding GFI1 motifs to simulated sequences containing one CRX motif at two sets of positions.

**Fig. S9:**
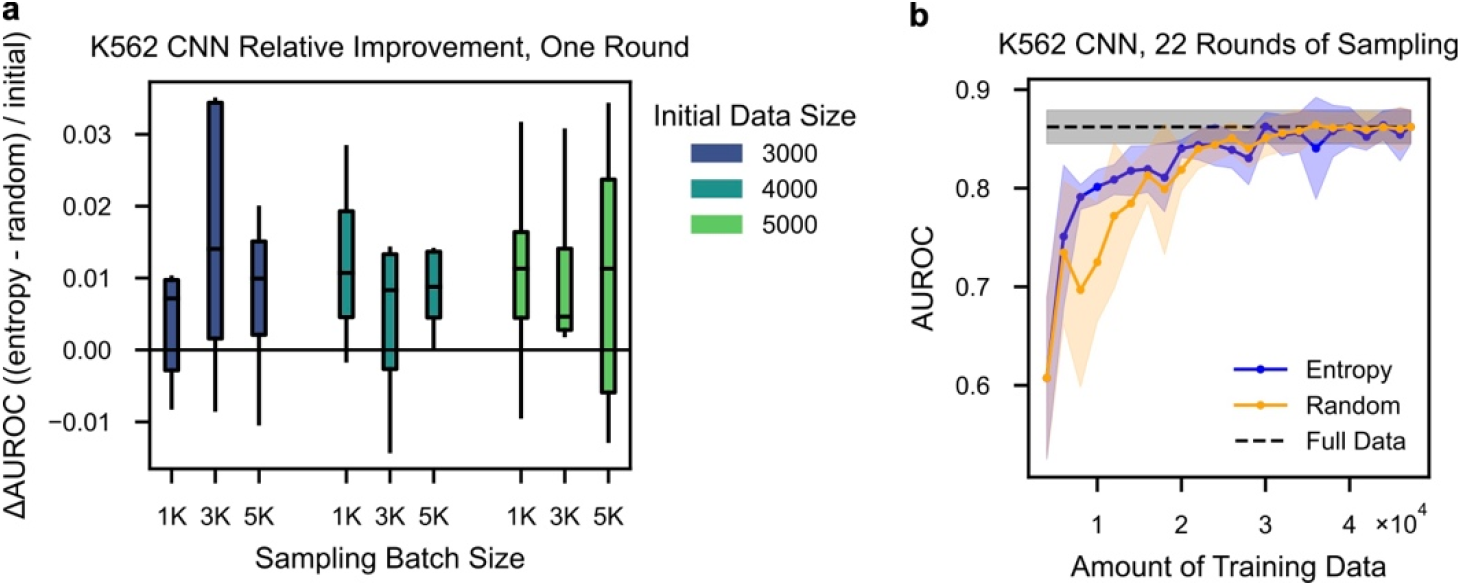
Additional benchmarking of active learning to K562 cells. **a**, Relative improvement of CNN for entropy sampling vs. random sampling compared to the initial performance, as measured by the AUROC on held-out test chromosomes. Each box represents 10-fold cross-validation for a set of starting conditions. **b**, Comparison of multiple rounds of entropy vs. random sampling for CNN performance as measured by AUROC. Lines denote the mean from 10-fold cross-validation, shaded areas denote 1 standard deviation, and black denotes CNN performance with the full dataset.

**Fig. S10:**
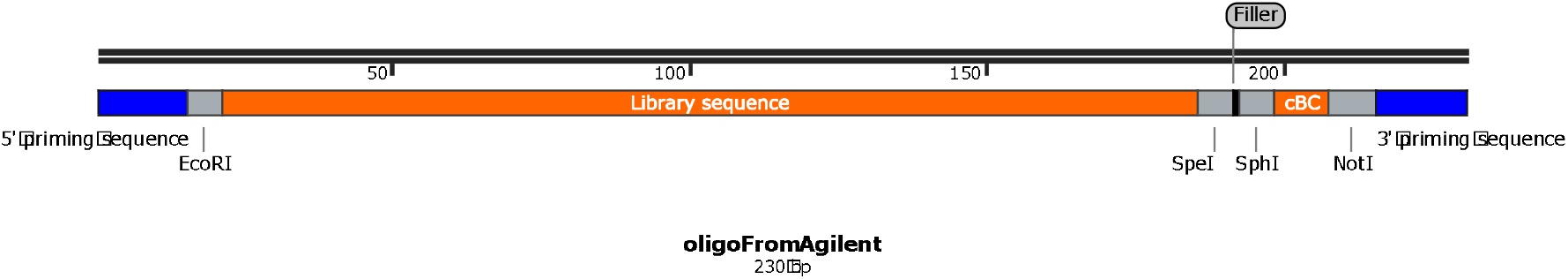
Snapgene schematic of MPRA library oligo design.

## SUPPLEMENTARY TABLES

**Supplementary Table 1: Library design experimental metadata.**

**Supplementary Table 2: mBC metadata.**

**Supplementary Table 3: Oligo amplification primer pair metadata.**

**Supplementary Table 4: Primers used in this study.**

**Supplementary Table 5: Within-library quality control statistics.**

**Supplementary Table 6: Block coordinates for the strong enhancer and inactive sequence pairs.**

**Supplementary Table 7: Annotations for swapping of blocks.** Uppercase letter denotes the block originates from the strong enhancer twin, lowercase denotes the inactive twin. A block letter followed by an ‘x’ indicates the region is scrambled, with the exact coordinates indicated by the numbers. All coordinates are on the zero-based interval [start, stop). Note that scrambles corresponding to motifs stretch 3bp into the flanks of adjacent blocks.

**Supplementary Table 8: Annotations for experiment sliding motifs along strong enhancers.**

